# The Role of TCF7L2 in Regulating Energy Metabolism in Thalamocortical Circuitry and its Broader Impact on Social Behavior

**DOI:** 10.1101/2025.11.14.684361

**Authors:** Andrzej Nagalski, Kamil Koziński, Suelen Baggio, Łukasz M. Szewczyk, Marcin A. Lipiec, Ben Hur M. Mussulini, Michael O. Gabriel, Joanna Bem, Ksenia Meyza, Anna Goncerzewicz, Ludwika Kondrakiewicz, Konrad Danielewski, Dariusz Gołowicz, Anna Kiryk, Urszula Wojda, Krzysztof Kazimierczuk, Ewelina Knapska, Agnieszka Chacińska, Marta B. Wiśniewska

## Abstract

Psychiatric and metabolic disorders often co-occur. While shared genetic factors and cellular dysfunctions are implicated, the underlying molecular mechanisms remain poorly understood. TCF7L2, a risk gene for type 2 diabetes and autism spectrum disorder, is highly expressed in the thalamus—a brain region extensively interconnected with the cortex, playing a key role in sensory processing, motor control, and behavioral regulation. Given its known role as a transcription factor regulating systemic energy metabolism, we explored its potential contribution to brain metabolism and behavior. To this end, we used a conditional knockout model with postnatal TCF7L2 loss in the thalamus and partial deficiency in the pancreas. *Tcf7l2* knockout mice exhibited social deficits and reduced motor habituation. In parallel, they also developed systemic glucose intolerance, modelling the psychiatric-metabolic comorbidity. Thalamic depletion of TCF7L2 resulted in elevated inhibitory phosphorylation of the pyruvate dehydrogenase —an enzymatic gatekeeper for pyruvate utilization in energy production—in the thalamus and cortex. This was accompanied by altered thalamic and cortical metabolism, characterized by reduced efficiency of pyruvate oxidation alongside enhanced oxidation of fatty acids and ketone bodies. Notably, a ketogenic diet alleviated metabolic dysregulation in the brain and normalized some social behaviors in knockout mice. These findings suggest that impaired energy metabolism in the thalamocortical circuitry may represent one of the pathogenic mechanisms underlying neuropsychiatric symptoms.

## Introduction

The prevalence of metabolic disorders is higher in individuals with psychiatric conditions. This association is particularly evident in autism spectrum disorder (ASD) and schizophrenia (SCZ), where the risk of developing metabolic conditions like type 2 diabetes or dyslipidemia is increased ^1–4^. These disorders may influence each other’s onset or progression and also appear to share genetic risk factors, dysregulated cell signaling pathways and mitochondrial dysfunctions, suggesting potential common origins ^5–8^. However, the molecular mechanisms linking metabolic and psychiatric disorders remain unclear.

Wnt/β-catenin signaling is one of the molecular pathways implicated in metabolic and psychiatric disorders ^9–11^. In particular, TCF7L2, a key effector transcription factor in the Wnt pathway, is predominantly expressed in metabolic organs and the brain. Variants of the *TCF7L2* gene are among the strongest genetic risk factors for type 2 diabetes ^12–14^. TCF7L2 is essential for peripheral metabolic homeostasis, regulating metabolic organ development ^15,16^, insulin production and secretion in pancreatic β-cells ^17,18^, metabolic adaptation in the liver ^19,20^, and lipid metabolism in adipose tissue ^21,22^. Beyond its role in diabetes, genetic variations in this gene have been associated with ASD and SCZ ^10,23–26^. Recently, mutations in *TCF7L2* have been identified as the cause of a rare neurodevelopmental disorder presenting with a spectrum of symptoms, including motor delay, intellectual disability, and features of autism, with variable expression across individual ^27^ (https://trndnetwork.org/trnd/signs-symptoms/).

In the brain, TCF7L2 is highly expressed in neurons within subcortical regions involved in sensory processing and behavioral response, where it promotes the expression of genes involved in connectivity development and maintenance of electrophysiological properties ^28–30^, oligodendrocyte precursors and young astrocytes ^24,31–34^. At the behavioral level, TCF7L2 regulates nicotine intake through habenular neurons ^35^, vocalization via periaqueductal grey neurons ^36^, and sociability through astrocytic functions ^24^. Nevertheless, the role of TCF7L2 in brain function and its contribution to the pathogenesis of ASD and SCZ remains far less understood than its role in metabolic organs.

The thalamus maintains the highest levels of TCF7L2 in the brain ^29^. This subcortical brain structure and its reciprocal cortical connections play a key role in modulating sensorimotor signaling and participate in various cognitive and behavioral functions ^37–40^. Neuroimaging studies repeatedly showed aberrant thalamocortical connectivity in ASD and SCZ individuals ^41–49^, suggesting thalamic dysfunctions in the pathogenesis of those disorders. Similarly, in mice, thalamocortical connection impairments were observed in classical models of ASD ^50,51^, and activity of specific thalamic nuclei was linked to social deficits ^52–58^. Furthermore, studies in genetic models confirmed a causal link between certain ASD risk genes, thalamic dysfunctions and ASD-relevant phenotypes, including attention deficits, seizures, sleep fragmentation, increased self-grooming, sensory disturbances and social deficits ^59–62^, with one study also reporting social deficits ^63^. They also highlighted the association between impaired neuronal inhibition linked to channelopathies and these phenotypes. However, experimental confirmation that thalamic dysfunction driven by ASD risk gene loss underlies core ASD manifestations— such as behavioral inflexibility and reduced sociability—remains limited, and other cellular mechanisms have yet to be explored.

Here, we demonstrate that postnatally induced deficiency of TCF7L2 in the thalamus affects social performance and motor flexibility in mice. We demonstrate that TCF7L2 plays a role in regulating energy metabolism in both the thalamus and cortex. Furthermore, we provide evidence that metabolic interventions can normalize brain metabolism and partially rescue behavioral deficits caused by *Tcf7l2* knockout, experimentally linking ASD with metabolic dysfunctions in the brain. Therefore, we propose that an imbalance in energy metabolism in the thalamocortical circuit can contribute to ASD pathomechanism.

## Results

*Cck*-driven Cre-mediated recombination of *Tcf7l2* removes TCF7L2 from the thalamus and partially from the pancreas, inducing a concurrent prediabetic state To examine the role of thalamic TCF7L2 in the development of social impairments and metabolic dysfunctions, we generated a conditional knockout of *Tcf7l2* in cells expressing the cholecystokinin (*Cck*) gene, whose expression overlaps with *Tcf7l2* expression in neurons in the thalamus. Crossing *Tcf7l2* floxed (*Tcf7l2^fl/fl^*) mice with *Cck^Cre^* animals results in the loss of TCF7L2 in most thalamic nuclei by postnatal day 14 (P14), as demonstrated in our previous study ^31^ (Fig. S1A–B, a schematic overview of the affected nuclei), and this loss was confirmed here at P60 (Fig. 1 A-B). We did not observe the knockout in other *Tcf7l2*-expressing brain regions, such as habenulae, dorsal midbrain ^31^ or hypothalamus (SFig. 2).

**Fig. 1.**
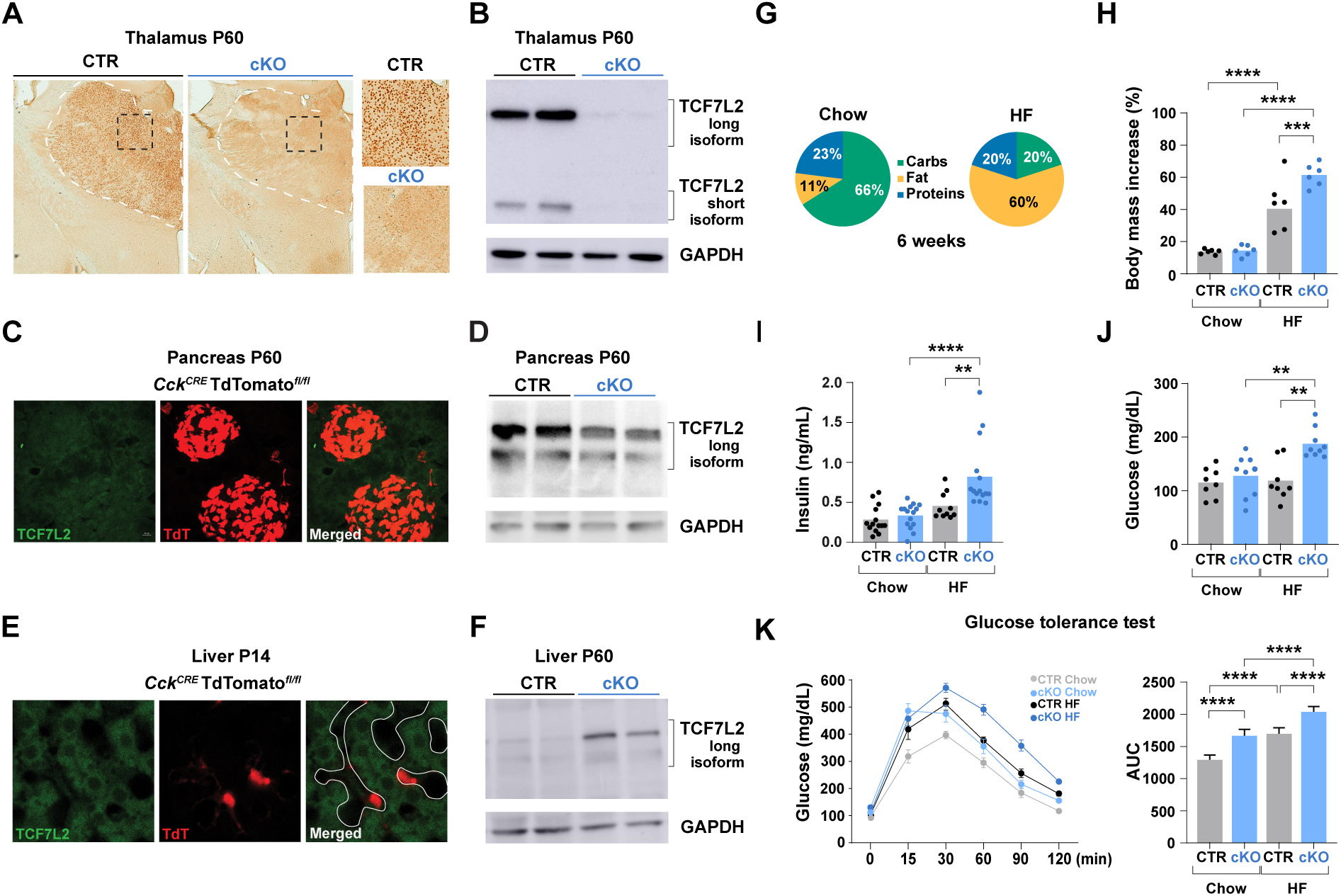
Cre-mediated recombination of *Tcf7l2* in *Cck*-expressing cells depletes TCF7L2 in the thalamus and partially in the pancreas, inducing a concurrent prediabetic state. **(A)** DAB immunohistochemical staining of TCF7L2 in coronal brain sections from P60 control (CTR) and *Cck^Cre^:Tcf7l2^fl/f^*^l^ (cKO) male mice; white dotted lines demarcate the thalamus. **(B)** Western blot analysis of TCF7L2 in the thalamus from P60 male mice; the upper band represents the long isoform, and the lower band the short isoform of TCF7L2. **(C)** Immunofluorescent staining of TCF7L2 (green) in the pancreas of the *Cck^Cre^:TdTomato^fl/fl^* reporter line at P60. **(D)** Western blot analysis of TCF7L2 in the pancreas from P60 male mice; only the long isoform of TCF7L2 is present. **(E)** Immunofluorescent staining of TCF7L2 (green) in the liver from the *Cck^Cre^:TdTomato^fl/f^*^l^ reporter line at P14. **(F)** Western blot analysis of TCF7L2 in the liver from P60 male mice. **(G)** Schematic of the dietary intervention showing the nutrient composition of the standard laboratory (Chow) and high-fat (HF) diet; only males were tested. **(H)** Percentage increase in body mass in male mice fed the chow or HF diet. **(I)** Fasting blood insulin levels. **(J)** Fasting blood glucose levels. **(K)** Glucose tolerance test (GTT; left panel); area under the curve (AUC) of GTT for the same groups (right panel); n = 6. Bars in H, I and J represent mean values, with dots indicating individual mice; dots in K represent mean values. Data in H, I, J, and K right panel were analyzed with two-way ANOVA followed by Tukey’s multiple comparison test. **p < 0.01, ***p < 0.001, ****p < 0.0001.

Given the metabolic focus of this research, it was important to first assess potential TCF7L2 deficiency in the *Cck^Cre^*:*Tcf7l2^fl/fl^* strain, hereinafter referred to as *Tcf7l2*-cKO, in the pancreas, liver and duodenum, where TCF7L2 plays a well-established role in regulating peripheral metabolism. To this end, we examined whether the *Cck* transgenic promoter drives *Cre* expression in the abovementiones organs using the *Cck^Cre^:TdTomato^fl/fl^* reporter line, and evaluated the impact of the conditional knockout on TCF7L2 expression.

We observed a TdTomato signal in numerous pancreatic cells, consistent with the moderate expression of *Cck* in the pancreas ^64,65^ which also stained positive for TCF7L2 (Fig. 1C). Histological features of these cells were characteristic of β-cells, suggesting that TCF7L2 could be removed from these cells. Western blot analysis further confirmed this, showing a decrease in TCF7L2 levels in the pancreas from *Tcf7l2*-cKO mice (Fig. 1D), indicating a partial pancreatic deficiency of TCF7L2 in our model.

In the liver, sparse TdTomato-positive cells were TCF7L2-negative (Fig. 1E). Their morphology and localization in the lumen of blood vessels indicated that they were not hepatocytes but either epithelial or Kupffer cells. This suggested a hepatocyte-specific gene knockout was unlikely. Unexpectedly, the level of TCF7L2, assessed by Western blot, increased in the liver (Fig. 1F), possibly as a secondary effect of the pancreatic knockout.

In the duodenum, TCF7L2-positive cells localized in crypts, while sparse TdTomato-positive cells were TCF7L2-negative (SFig. 3), excluding *Tcf7l2* knockout in this organ. The localization of TdTomato-positive cells was consistent with the localization of *Cck*-expressing enteroendocrine cells ^66^.

Decreased TCF7L2 levels in the pancreas and increased levels in the liver suggested a complex misregulation of *Tcf7l2* expression in our model, potentially contributing to metabolic dysfunction. To test this, we assessed body weight, glucose, and insulin levels in control and *Tcf7l2*-cKO mice fed either a standard chow (66% energy from carbohydrates) or high-fat (HF, 60% energy from fat) diets for six weeks (Fig. 1G). On the chow diet, control and *Tcf7l2*-cKO mice gained weight similarly (Fig. 1H). Fasting insulin and glucose levels were also comparable (Fig. 1I-J), but *Tcf7l2*-cKO mice showed impaired glucose clearance (Fig. 1K). Under the HF diet, metabolic differences became more pronounced: *Tcf7l2*-cKO mice gained significantly more weight (Fig. 1H), exhibited elevated fasting insulin and glucose (Fig. 1I-J), and showed exacerbated glucose intolerance (Fig. 1K).

Our approach effectively eliminated TCF7L2 from most thalamic neurons and partially from pancreatic β-cells. Consequently, the transgenic mice displayed a prediabetic state alongside potential thalamic impairments, rendering them suitable for investigating the comorbidity of thalamus-related psychiatric disorders and metabolic dysfunctions.

### *Cck^Cre^*:*Tcf7l2^fl/fl^* mice show no evidence of impairment in basic locomotor parameters, anxiety-like behavior, or major cognitive deficits

Patients with mutations in the *TCF7L2* gene exhibit impaired motor function, increased anxiety, cognitive disabilities and difficulties with social interaction ^27^. To determine if postnatal thalamic ablation of TCF7L2 could lead to similar behaviors, we conducted a series of behavioral tests in *Tcf7l2*-cKO mice. Both adult males and females were included unless otherwise noted.

We started with assessing basic locomotor and sensorimotor performance. Locomotor functions were evaluated in a dimly lit, non-anxiogenic enclosure enriched with objects (Fig. 2A). Under these conditions, CTR and *Tcf7l2*-cKO mice displayed comparable immobility times, average and maximal speeds, and angular velocity, in both sexes (Fig. 2B-D). We then assessed olfactory function using a buried food-seeking test (Fig. 2E). Latency to retrieve the food did not differ between genotypes (Fig. 2F). These results showed intact locomotor and sensorimotor functions *Tcf7l2*-cKO mice under low-demand conditions.

**Fig. 2.**
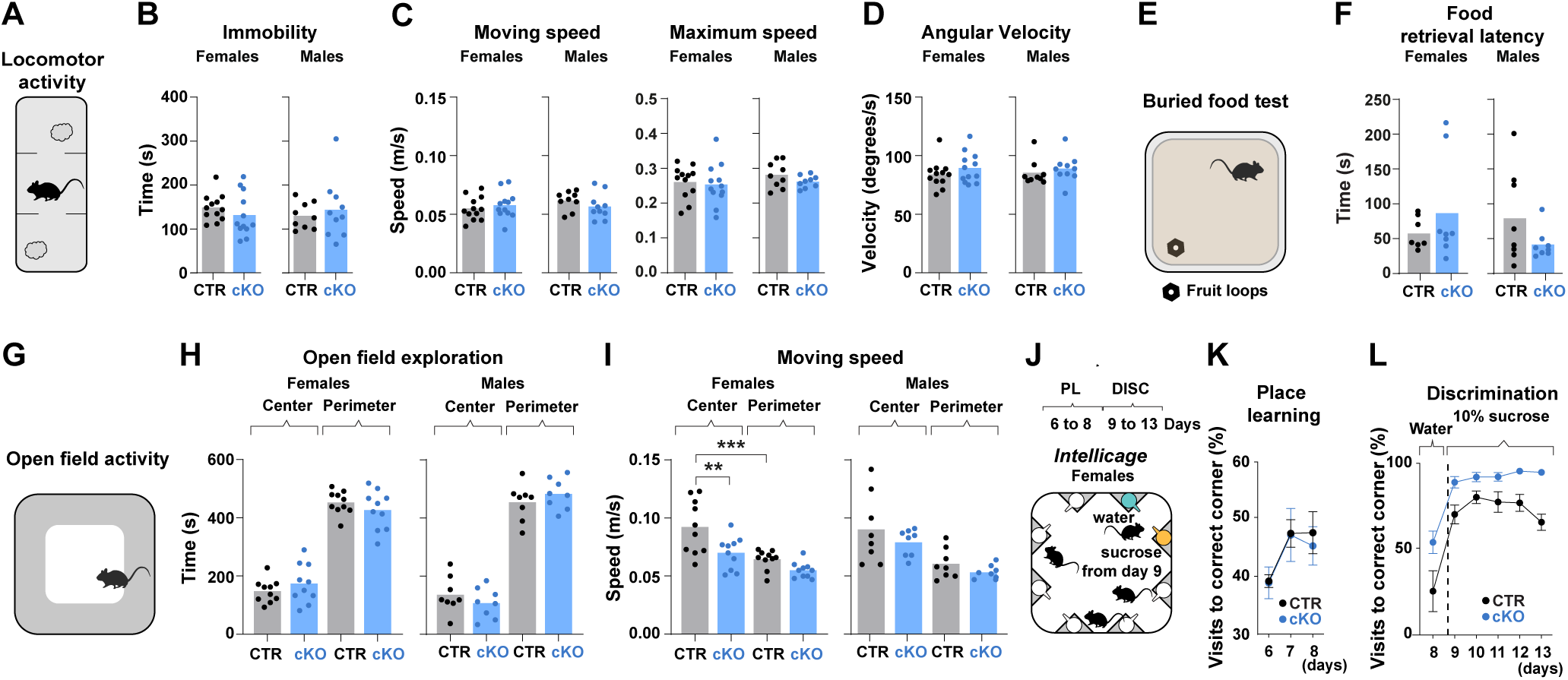
TCF7L2 depletion in the thalamus does not cause apparent alterations in locomotor, anxiety-related, or cognitive performance. **(A)** Schematic of a dimly light enclosure enriched with objects; both sexes were tested. Data were recorded for 10 minutes. **(B)** Immobility time of control (CTR) and *Cck^Cre^:Tcf7l2^fl/f^*^l^ (cKO) mice. **(C)** Average (left panel) and maximum (right panel) speed when moving. **(D)** Average angular velocity. **(E)** Schematic of the buried food test used to assess olfactory function; both sexes were tested. **(F)** Latency to retrieve hidden food. **(G)** Schematic of the open field test; both sexes were tested. **(H)** Time spent in the center or the perimeter of the open field. **(I)** Mobility speed in the center or the perimeter of the open field. **(J)** Schematic of the *Intellicage* setup used to assess learning and motivation. During the nose-poke adaptation phase (days 4-5), mice accessed water in all corners and learned to nose-poke; in the place learning phase (PL, days 6-8), each mouse was assigned a specific corner, while the remaining corners were inaccessible, in the cue discrimination (reward-based learning) phase (DISC, days 9-13), one corner bottle was replaced with 10% sucrose. CTR n = 23 and cKO n = 18 in two independent cohorts for each genotype; only females were tested; results in K-L represent data from the dark phase. **(K)** Percentage of visits to the correct (assigned) corner relative to total visits to all corners, days 6–8. **(L)** Percentage of visits to the sucrose-containing bottle relative to total visits to the correct corner, days 9–12. Bars in graphs B, C, D, F, H, and I represent mean values, with dots indicating individual mice; dots in K and L represent mean values. Data in B, C, D, and F were analyzed using an unpaired T-test; data in H, I, K, and L were analyzed using two-way ANOVA followed by Tukey’s multiple comparison test.

Next, to assess risk-taking and anxiety-related behaviors, we used the light–dark box (SFig. 4A) and open-field (Fig. 2G) apparatuses. In the light–dark box, there were no genotype differences in the time spent in the light zone, latency to enter the dark zone, or the number of transitions between compartments (SFig. 4B–D). Similarly, no significant differences were observed between control and *Tcf7l2*-cKO mice in the time spent in the center of the open field (Fig. 2H). Analysis of locomotor activity, however, during the open-field test revealed a genotype-dependent difference. While mice of both genotypes moved at similar speeds along the perimeter (Fig. 2I), control mice increased their locomotor activity in the center, moving faster than at the periphery, with this effect reaching statistical significance in females. In contrast, *Tcf7l2*-cKO mice lacked this modulation, suggesting impaired context-dependent locomotor adjustment. These results indicated that postnatal *Tcf7l2* knockout in thalamic neurons does not induce anxiety-like behaviors but may alter locomotor adaptability.

We then assessed cognitive performance using an automated *IntelliCage* system, which tracked individual mice as they visited water bottles in corners within a group-housed setting via implanted transponders (Fig. 2J). Due to aggressive behaviors between males within a group, we tested cohorts composed only of females. The testing protocol consisted of place learning and reward-based discrimination learning tasks, preceded by adaptation phases. During the place learning phase, each mouse learned to access ts individually assigned corner, a task that required spatial associative learning. Females of both genotypes performed equally well, as measured by an increasing percentage of correct visits (Fig. 2K). During the reward-based learning phase, one of the two bottles in the assigned corner was filled with sweetened water. Mice from both groups rapidly developed a preference for the rewarding bottle (Fig. 2L), with no significant differences in performance. These results suggested no overt deficits in spatial or reward-based learning and motivation in *Tcf7l2*-cKO mice under these conditions.

### Postnatal thalamic knockout of *Tcf7l2* leads to social deficits and reduced motor habituation

In the next part of the behavioral testing, we focused on social behaviors, using the classic three-chamber test (interest in social stimulus), the *IntelliCage* system (group preference for a corner during the simple adaptation phase), and the *EcoHAB* setup (dynamic and passive interactions within a group).

We began with evaluating the mice’s interest in social over non-social interactions in the three-chamber test (A). In the apparatus, mice were exposed to an unfamiliar mouse under one cup (social stimulus) and a new inanimate object (non-social stimulus) under another. Control mice spent significantly more time interacting with the mouse than with the object (Fig. 3B). Similarly, *Tcf7l2*-cKO males displayed normal social interest. In contrast, knockout females exhibited a marked impairment in social preference, as evidenced by a significantly lower discrimination index.

**Fig. 3.**
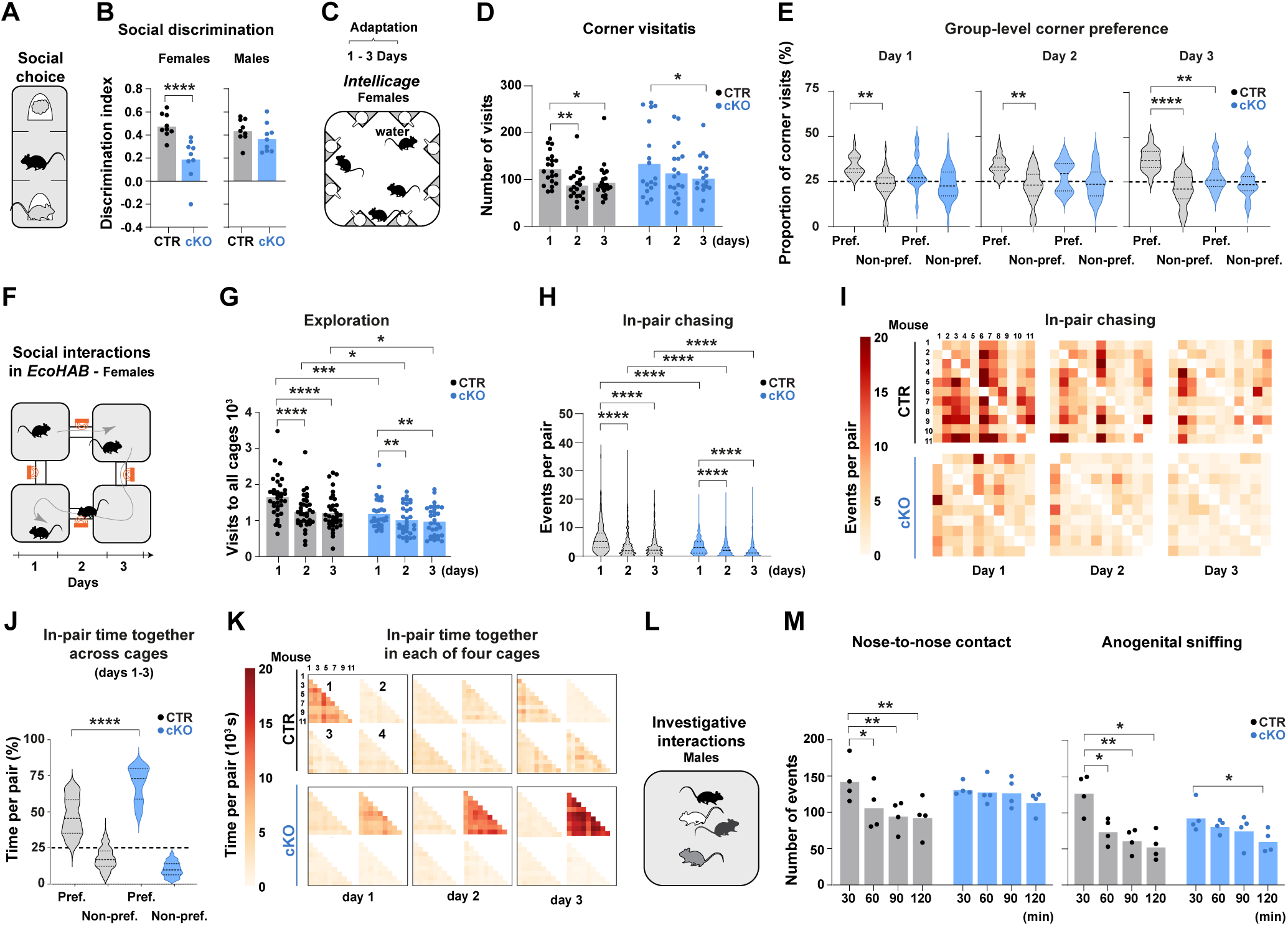
Postnatal thalamic knockout of *Tcf7l2* leads to social deficits. **(A)** Schematic of the three-chamber social interest test; a mouse was presented with an unfamiliar mouse and an object; both sexes were tested. **(B)** Preference for a mouse over an object, calculated for control (CTR) and *Cck^Cre^:Tcf7l2^fl/f^*^l^ (cKO) mice as the discrimination index. **(C)** Schematic of the *Intellicage* setup showing water access in four corners during the simple adaptation phase (days 1-3). CTR n = 23 and cKO n = 18 in two independent cohorts for each genotype; only females were tested; results in D-E represent data from the dark phase. **(D)** Number of visits to all corners. **(E)** Percentage of each mouse’s visits to the group’s most preferred corner (Pref.) or non-preferred corners (Non-pref.), relative to its total number of visits on each of three days; an alternative visualization of the same result is provided as a bar plot in SFig. 5B. **(F)** Schematic of the *EcoHAB* setup used to assess social behavior. The setup consisted of four cages connected by tunnels with antennas to track mouse movements. CTR n = 35 and cKO n = 32 in three independent cohorts for each genotype; only females were tested; results in G-I represent data from the dark phase. **(G)** Exploratory behavior, measured as the total number of visits to all cages. **(H)** In-pair chasing events, measured as one mouse trailing another through a tunnel for each mouse pair; an alternative visualization of the same result is provided as a bar plot in SFig. 5E. **(I)** Representative heatmaps (right panel) showing the number of in-pair chasing events for one control and one cKO cohort; columns represent chasing mice, rows represent chased mice; the scale was limited to 20 for better visualization of the result. **(J)** Percentage of in-pair time spent spent together in the group’s most preferred cage (Pref.) or non-preferred cages (Non-pref.), relative to total time spent together, calculated for each pair across days 1–3. **(K)** Representative heatmaps, from the same cohort as in panel I, showing in-pair time spent together for one control and one cKO cohort; the scale was limited to 20 for better visualization of the result. **(L)** Schematic of the social interaction test; four unfamiliar mice were placed in a cage for two hours; only males were tested. **(M)** Number of nose-to-nose (right panel) and anogenital sniffing (right panel) events. Bars in D, G, H, and M represent mean values, with dots indicating individual mice; violin plots in E, H, and J represent the distribution of individual data points, with the central line indicating the median and dashed lines indicating the interquartile range. Data in B were analyzed using an unpaired T-test; data in E and J were analyzed using two-way ANOVA followed by Tukey’s multiple comparison test; data in D, G, H, and M were analyzed using repeated measures two-way ANOVA followed by Tukey’s multiple comparison test. *p < 0.05, **p < 0.01, ***p < 0.001.

Next, we used the *IntelliCage* system, in which water bottles were located in four corners, over a three-day adaptation period (Fig. 3C) to examine whether patterns of corners’ visitations reflect social coordination and differ between genotypes. Females of both genotypes visited the *IntelliCage* corners at overall similar levels throughout the adaptation phase (Fig. 3D). Control females showed a reduction in corner visitation already on day 2, whereas a similar decrease in *Tcf7l2*-cKO females was observed on day 3. This temporal shift may indicate a delay in locomotor habituation in knockout mice.

Despite decreased visitation levels, control females exhibited a statistically significant emergence of a shared corner preference. By day 3, nearly 40% of all visits were directed toward a single common corner (Fig. 3E and SFig. 5A-B). Unlike control animals, *Tcf7l2*-cKO females failed to form such a group-level bias, suggesting decreased sensitivity to social cues.

Subsequently, to further investigate how female *Tcf7l2*-cKO mice interact within a group, we employed the *EcoHAB* system, which has been previously used to test mouse models of autism ^67^. In this naturalistic setup, female lived together for several days in a four-cage setup connected by tunnels (Fig. 3F). Each mouse was individually identified with an implanted transponder, and its movements between cages were continuously tracked.

We first analyzed exploratory behavior by counting the number of visits to all cages in three consecutive days (dark phases). On day 1, *Tcf7l2*-cKO mice made significantly fewer passages than control mice (Fig. 3G). In both groups, exploration declined on days 2 and 3; however, the reduction was smaller in *Tcf7l2*-cKO mice, with a decrease of approximately 25% in controls and 15% in *Tcf7l2*-cKO mice (SFig. 5C-D), suggesting reduced motor habituation, similar to *Intellicage*.

To assess dynamic social interactions during this period, we counted the number of times one mouse chased another during passages between cages through the connecting tunnels, for each pair. Control mice showed high in-pair chasing on average on the first day, with about a 40% decline in the subsequent days (Fig. 3H, violin plot; SFig. 5E, bar plot). This pattern was expected, as typical social behavior involves an initial increase in interactions to establish a social structure, followed by a decrease as the hierarchy forms. In contrast, *Tcf7l2-*cKO mice exhibited significantly reduced in-pair chasing levels. The effect was particularly pronounced on day 1, when in-pair-chasings were nearly twofold less frequent compared to control cohort.

Moreover, heatmaps of chasing interactions and their directionality (Fig. 3I) revealed individually specific patterns of social engagement: some mice were predominantly chasing, others were mostly being chased, some exhibited a similar level of chasing and being chased, and others showed minimal involvement in either behavior. Variability in in-pair chasing behavior was more pronounced in control mice, a difference between genotypes that was statistically confirmed by Levene’s test (p < 10⁻⁸, p < 0.001, and p = 0.02 for control–*Tcf7l2*-cKO comparisons on days 1, 2, and 3, respectively). Pearson’s correlation analyses across consecutive days revealed that individual chasing patterns were stable across consecutive days (SFig. 5F), indicating that the observed variation reflected consistent behavioral tendencies rather than random fluctuations. Notably, this temporal stability between day 1 and 2 was greater in control mice than in *Tcf7l2*-cKO mice. Together, these findings indicated that genotype-dependent differences in chasing affected not only the overall level, but also the structure of social engagement.

Our previous work showed that chasing behavior serves to manifest dominance ^68^, and the results described above further indicated that chasing captures stable, individual-specific patterns of social organization within the group. Nevertheless, to rule out the possibility that reduced chasing in *Tcf7l2*-cKO mice is a consequence of lower exploratory drive, we normalized the data by calculating the proportion of socially motivated passages (i.e., chasing) relative to the total exploratory activity of a mouse (SFig. 5G). This proportion was significantly lower in *Tcf7l2*-cKO mice, supporting reduced active social engagement. In addition, Pearson’s correlation analysis revealed that exploration correlated weakly or not at all with being chased, with no statistical significance in most cases (SFig. 6A-G) and moderately with chasing (SFig. 6H-M), indicating that these two components of social behavior relate differently to exploration.

As an additional measure of social behavior, we calculated, for each mouse pair, the proportion of their total time spent together that occurred in the cage most preferred by the group (i.e., the cage in which mouse pairs spent the most of their time together on average). All cohorts exhibited some degree of consensus in cage preference, with *Tcf7l2*-cKO mice showing an extreme pattern—spending nearly all of their in-pair time together in a single cage (Fig. 3J-K). These patterns suggested reduced flexibility in how *Tcf7l2*-cKO mice interact with their social environment.

Due to aggressive behavior, males were not tested in *EcoHAB*, as was the case in *IntelliCage*. Instead, groups of four unfamiliar males were placed together in a shared cage for two hours (Fig. 3L), during which spontaneous direct social interactions were recorded, including investigative behaviors such as nose-to-nose and anogenital sniffing. During the first 30 minutes, *Tcf7l2*-cKO mice exhibited a similar number of nose-to-nose contacts but fewer occurrences of anogenital sniffing compared to controls (Fig. 3M), indicating reduced social investigative behaviors. Notably, while control mice showed a significant reduction in interactions during the subsequent 30-minute intervals (by 30-40%), *Tcf7l2*-cKO mice maintained a steady level of interactions throughout the entire test period, suggesting a lack of short-term social habituation.

Taken together, the results from *IntelliCage*, *EcoHAB*, the three-chamber test and the investigative interaction test suggested that *Tcf7l2*-cKO mice have a reduced capacity to adapt their behavior to changing social and environmental conditions.

### TCF7L2 regulates metabolic genes and metabolic pathways in the thalamus

Given TCF7L2’s established role in regulating metabolic gene expression in the pancreas ^69^ and liver ^20^, we hypothesized that TCF7L2 similarly influences metabolic gene expression in the thalamus. To test this, we analyzed our previous RNA-seq data, focusing on metabolism-associated genes, and integrated the results with our TCF7L2 ChIP-seq data from the same region ^31^, thalamic expression enrichment from the Allen Brain Atlas (available from mouse.brain-map.org) ^70^ and cell-type enrichment based on published single-cell RNA-seq data from the thalamus ^71^.

We identified 49 metabolism-related differentially expressed genes (DEGs) in the thalamus of knockout mice compared with control mice, of which 30 were downregulated and 19 were upregulated (Fig. 4A). More than 30% of these genes were characterized by the presence of TCF7L2 ChIP-seq peaks, indicating direct regulation by this factor. The majority were neuron-enriched. Based on their cellular functions, we grouped the genes into categories related to energy metabolism. Within these categories, we identified two particularly interesting gene sets.

**Fig. 4.**
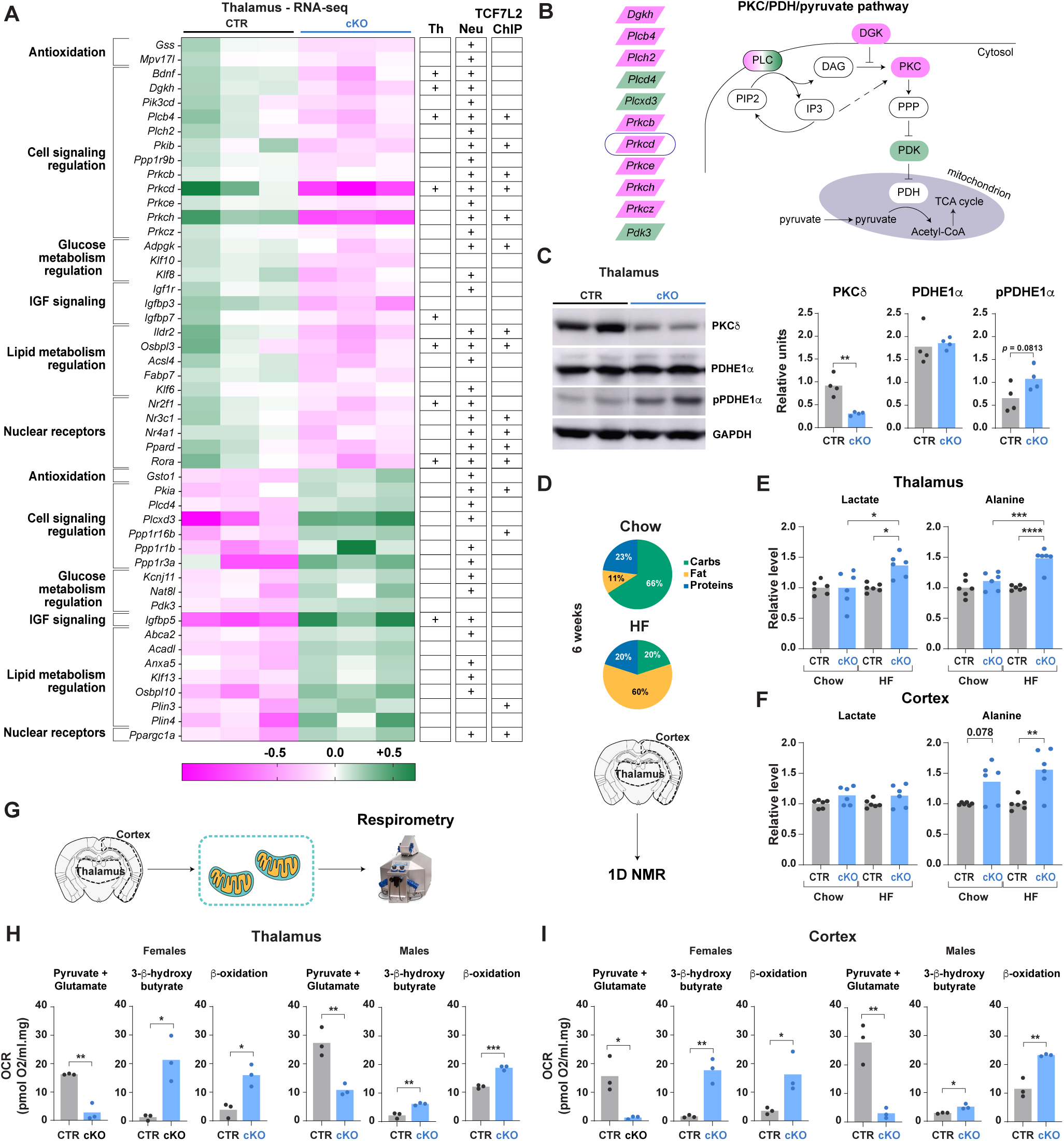
Postnatal thalamic knockout of *Tcf7l2* influences metabolic gene expression and pyruvate dehydrogenase activity, affecting the utilization of pyruvate in the thalamus and cortex. **(A)** Upregulated and downregulated genes in the thalamic region of *Cck^Cre^:Tcf7l2^fl/f^*^l^ (cKO) mice compared to control (CTR) mice. Genes were manually curated into the functional categories indicated on the left. Log2-transformed expression fold changes (cKO/CTR), centered to the median of each row, are displayed using a magenta–green color scale, each column represents an independent biological replicate. Right panel: thalamic enrichment (Th) is based on the Allen Brain Atlas; neuronal enrichment (Neu) is based on a whole-brain single-cell RNA-seq database, and positive TCF7L2 ChIP signal is based on our published results. **(B)** Schematic of a hypothetical signaling pathway affected by the conditional knockout of *Tcf7l2*. **(C)** Western blot analysis of PKCδ, PDHE1α, and phospho-PDHE1α in the thalamus of male mice. **(D)** Schematic of the dietary intervention showing the nutrient composition of the standard laboratory (Chow) and high-fat (HF) diet, followed by brain sectioning for 1D NMR analysis; only males were tested. **(E-F)** Levels of lactate and alanine in the thalamus and cortex of mice fed the Chow or HF diet. **(G)** Schematic of brain sectioning and subsequent cell permeabilization for high-resolution respirometry analysis; both sexes were tested. **(H-I)** Oxygen consumption rates (OCR) of permeabilized tissue membranes isolated from the thalamus and cortex of female and male mice. Data were obtained by sequentially adding fatty acids, 3-β-hydroxybutyrate, and pyruvate + glutamate to the Oxygraph-2k chamber. Bars in graphs represent mean values, with dots indicating individual mice. Data in C, E, F, H, and I were analyzed using an unpaired T*-*test. *p < 0.05, **p < 0.01, ***p < 0.001, ****p < 0.0001.

The first group comprised DEGs involved in PPAR/fatty acids signaling pathways, including nuclear receptor transcription factors *Ppard, Rora*, and *Ppargc1a*, as well as lipid metabolism-related genes *Plin3*, *Plin4*, *Acsl4*, *Acadl, Igfbp3,* and *Igfbp5*. The presence of direct TCF7L2 binding sites at *Ppard* and *Rora*—but not at the other genes—suggests that TCF7L2 may regulate lipid metabolism indirectly through these nuclear receptors.

The second set of DEGs was associated with the protein kinase C (PKC) pathway, known for regulating energy metabolism and glucose homeostasis ^72^. (Fig. 4B). This group included phospholipases *Plcb4*, *Plch2* and *Plcd*, a diacylglycerol kinase *D*g*kh*, protein kinases *Prkcb*, *Prkcd* (PKCδ), *Prkce*, *Prkch* and *Prkc,* and pyruvate dehydrogenase *Pdk3*. TCF7L2-ChIP-seq peaks were mapped to *Plcb4*, *Prkcb*, *Prkcd* and *Prkch*, suggesting direct regulation of these genes by TCF7L2.

We hypothesize that the PDHc/pyruvate pathway might be disrupted in the thalamus in the*Tcf7l2*-cKO mice. This was based on *vitro* evidence that PKCδ can signal to the pyruvate dehydrogenase complex (PDHc) which controls pyruvate utilization in the tricarboxylic acid cycle (TCA) ^73^, and on data showing elevated serum pyruvate in PKCδ-deficient mice ^74^. To investigate our prediction, we compared PKCδ levels and PDHE1α phosphorylation between control and knockout mice using Western blot. As expected from its transcript level, PKCδ was significantly downregulated in the thalamus of *Tcf7l2*-cKO mice (Fig. 4C). Consistent with the predicted signaling pathway, PDHE1α phosphorylation increased, while the total level of PDHE1α remained unchanged.

In summary, our findings indicate that TCF7L2 regulates the activity of at least two critical metabolic groups of pathways: the PPARD pathways, through direct control of *Ppard* gene expression, and the PKC/PDHc pathways by directly modulating the expression of genes encoding PKCδ and other proteins associated with the pathway. This regulatory role suggested that TCF7L2 may shape the energy metabolism profile in the thalamus.

### Deficiency of TCF7L2 alters the profile of energy metabolism in the thalamus and cortex

To examine whether the thalamic loss of TCF7L2 affects brain metabolism at the functional level, we began by measuring soluble metabolites associated with glycolysis and the TCA cycle using ^1^H-NMR spectroscopy. This approach provided a broad overview of over a dozen water-soluble metabolites, most of which were linked to the TCA cycle (SFig. 7A). We analyzed both thalamic and cortical samples, assuming that metabolic impairments in the thalamus might impact the cortex due to the high presence of thalamus-derived presynaptic proteins and mitochondria there. Samples were collected from mice maintained on either the standard chow or HF diet, the latter included to expose potential metabolic defects under metabolic stress (Fig. 4D).

In knockout mice on the chow diet, levels of the measured metabolites were comparable to those in controls (Fig. 4E-F and SFig. 7A-D). However, when fed the HF diet, *Tcf7l2*-cKO mice exhibited significantly elevated lactate and alanine—both pyruvate-related metabolites—in the thalamus (Fig. 4E), and increased alanine in the cortex (Fig. 4F). These elevated levels may reflect altered pyruvate utilization, prompting us to further explore brain mitochondrial energy metabolism in these mice.

To directly assess mitochondrial oxidative capacity, we performed high-resolution respirometry on isolated thalamic and cortical cells (Fig. 4G), enabling us to evaluate intrinsic mitochondrial function independent of potential in vivo compensatory mechanisms. This analysis was conducted in both sexes.

Within mitochondria, pyruvate, ketone bodies and fatty acids are converted to acetyl CoA via distinct pathways: oxidative decarboxylation in the PDHc, oxidation by β-hydroxybutyrate dehydrogenase, and β-oxidation, respectively ^75^. To assess the efficiency of these pathways, we supplied our samples with palmitoylcarnitine in the presence of malate and non-limiting levels of ADP, octanoylcarnitine, 3-β-hydroxybutyrate, followed by pyruvate and glutamate. In the mitochondria from *Tcf7l2*-cKO mice, pyruvate oxidation was markedly reduced both in the thalamus and cortex of females and males (Fig. 4H-I). These results were consistent with reduced PDHc activity. In contrast, the oxidation of 3-β-hydroxybutyrate and fatty acid β-oxidation were significantly elevated in the same samples.

To determine whether these metabolic alterations were brain region-specific, we measured substrate oxidation rates in cerebellar mitochondria. We chose the cerebellum because it lacks input from the thalamus. No differences were detected in the efficiency of pyruvate oxidation, 3-β-hydroxybutyrate oxidation, or β-oxidation between knockout and control mice (SFig. 8). This finding suggests that the metabolic impairments observed in the cortex and thalamus of *Tcf7l2*-cKO mice were brain-region-specific and were unlikely to be secondary effects of peripheral metabolic changes.

Altogether, these findings demonstrate that the deficiency of thalamic TCF7L2 alters the energy metabolism profile in the thalamocortical circuitry, affecting the rates of energy production from different sources.

### A ketogenic diet mitigates brain metabolic and social deficits induced by thalamic *Tcf7l2* knockout

Given the altered brain metabolism and reduced social preference observed in *Tcf7l2-*cKO, we sought to investigate if a ketogenic diet, which has shown promise in treating individuals with ASD ^76–78^ and is recognized for its potential to modify metabolic pathways ^79^, could improve their social performance. To test this, we administered a ketogenic diet (94% of energy from fat) for six weeks (Fig. 5A) and assessed its impact on systemic metabolism, brain metabolism and social performance.

**Fig. 5.**
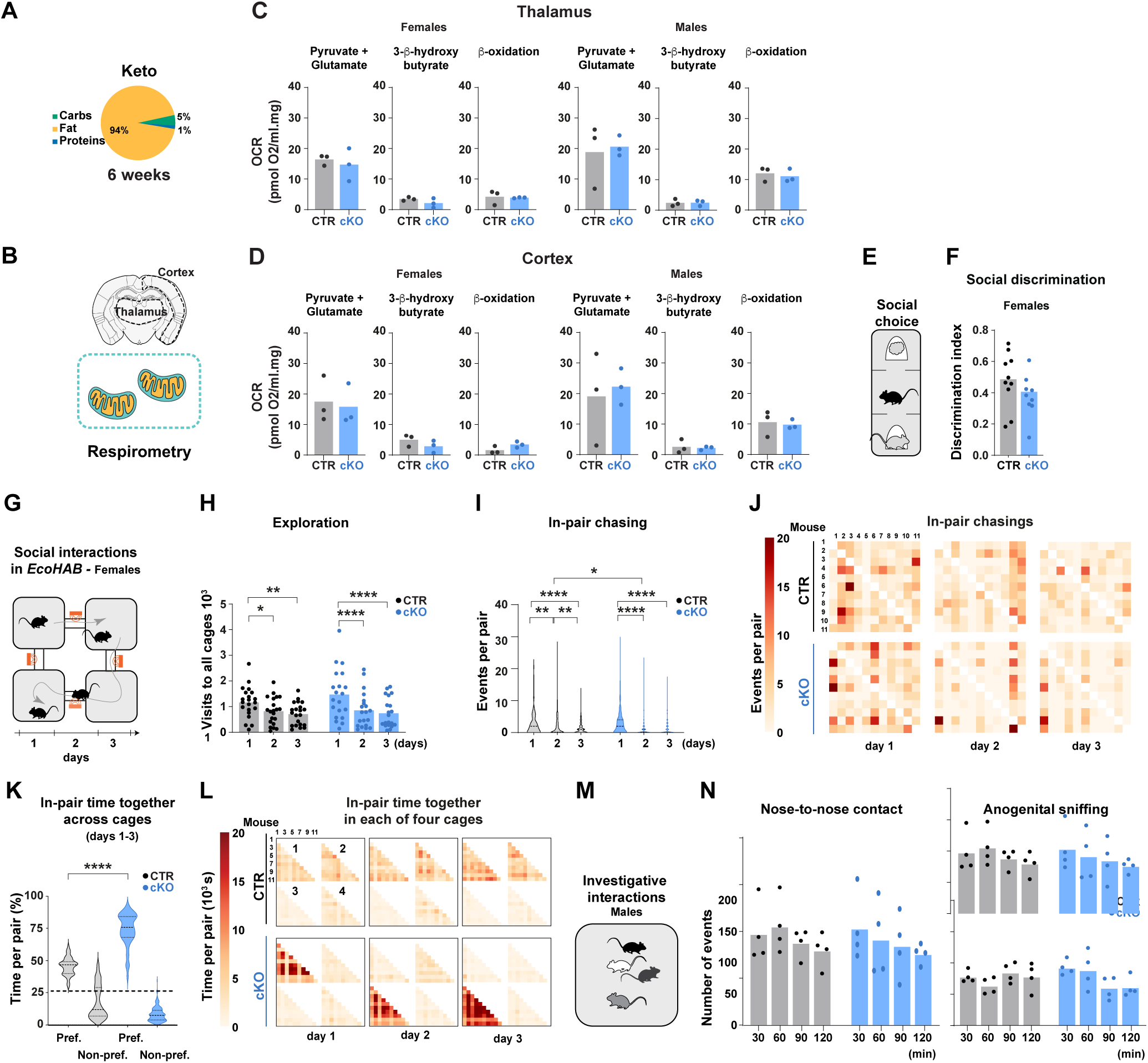
Ketogenic diet ameliorates brain metabolic and social deficits induced by postnatal thalamic knockout of *Tcf7l2*. **(A)** Schematic of the dietary intervention showing the nutrient composition of the ketogenic (Keto) diet. **(B)** Brain regions analyzed by high-resolution respirometry in permeabilized cells. **(C-D)** Oxygen consumption rates (OCR) of permeabilized tissue membranes isolated from the thalamus or cortex of female and male control (CTR) and *Cck^Cre^*:*Tcf7l2 ^fl/fl^* (cKO) mice fed either the Chow or Keto diet. Data was obtained by sequentially adding fatty acids, 3-β-hydroxybutyrate, and pyruvate + glutamate to the Oxygraph-2k chamber. **(E)** Schematic of the three-chamber social interest test, in which a mouse is presented with an unfamiliar mouse and an object; only females were tested. **(F)** Preference for a mouse over an object, calculated as the discrimination index. **(G)** Schematic of the *EcoHAB* setup used to assess social behavior. The setup consisted of four cages connected by tunnels with antennas to track mouse movements. CTR n = 21 and cKO n = 20 in two independent cohorts for each genotype; only females were tested; results in H-J represent data from the dark phase, days 1-3. **(H)** Exploratory behavior, measured as the total number of visits to all cages. **(I)** In-pair chasing events, measured as one mouse trailing another through a tunnel for each mouse pair; an alternative visualization of the same result is provided as a bar plot in SFig. 11D. **(J)** Representative heatmaps showing the number of in-pair chasing events from one CTR and one cKO cohort; columns represent chasing mice, rows represent chased mice; the scale was limited to 20 for better visualization of the results. **(K)** Percentage of in-pair time spent together in the group’s most preferred cage (Pref.) or non-preferred cages (Non-pref.), relative to total time spent together, calculated for each pair across days 1–3. **(L)** Representative heatmaps, from the same cohort as in panel J, showing in-pair time spent together for one control and one cKO cohort; the scale was limited to 20 for better visualization of the result. **(M)** Schematic of the social interaction test; four unfamiliar mice were placed in a cage for two hours; only males were tested. **(N)** Number of nose-to-nose (right panel) and anogenital sniffing (right panel) events. Bars in C, D, F, H, and N represent mean values, with dots indicating individual mice; violin plots in I and K represent the distribution of individual data points, with the central line indicating the median and dashed lines indicating the interquartile range. Data in C, D, and F were analyzed using an unpaired T-test; data in H, I, and N were analyzed using repeated measures two-way ANOVA followed by Tukey’s multiple comparison test; data in K were analyzed using two-way ANOVA followed by Tukey’s multiple comparison test. *p < 0.05, **p < 0.01, ***p < 0.001, ****p < 0.0001.

The ketogenic diet halted weight gain in control mice but did not affect weight gain in *Tcf7l2*-cKO mice compared to the chow diet (SFig. 9A-B). It elevated fasting insulin levels only in *Tcf7l2*-cKO mice (SFig. 9C). Although fasting glucose levels remained unaffected (SFig. 9D), glucose tolerance decreased in both genotypes compared to the chow diet, with a more pronounced effect observed in knockout mice (SFig. 9E). These changes indicated the emergence of insulin resistance, comparable to metabolic effects observed with HF diet exposure. Collectively, these results indicated the ketogenic diet adversely affected systemic metabolism homeostasis in both genotypes.

Next, we examined the impact of the ketogenic diet on energy metabolism in the thalamus and cortex using high resolution respirometry (Fig. 5B). Notably, the ketogenic diet normalized respiration rates in both brain regions in *Tcf7l2*-cKO mice, in both females and males (Fig. 5C-D). In particular, pyruvate oxidation levels in knockout mice increased to levels comparabe to those of control animals. Thus, surprisingly, while systemic metabolism deteriorated, the ketogenic diet led to improvements in brain energy metabolism.

Finally, we assessed the effects of ketogenic diet on social behavior and locomotor habituation using the three-chamber test and the *EcoHAB* system (in females), as well as a spontaneous social direct interaction test in a group of unfamiliar mice (in males).

In the three-chamber test (Fig. 5E), the ketogenic diet did not affect social preference in control females but increased interest in the unfamiliar mouse in *Tcf7l2*-KO females, restoring their preference to control levels (Fig. 5F).

In *EcoHAB* (Fig. 5G), the ketogenic diet reduced exploratory behavior in control mice while increasing it in *Tcf7l2*-cKO mice (Fig. 5H), thereby eliminating the genotype-related difference observed under standard diet conditions (Fig. 5H; compare with Fig. 3G). Simultaneously, the ketogenic diet restored locomotor habituation in *Tcf7l2*-cKO mice (SFig. 10A-C; compare with SFig. 5D), as indicated by an approximately 50% reduction in cage visits—an even greater decrease than that observed in controls (30%). Similarly, in-pair chasing interactions declined in control mice on day 1, reaching levels observed in *Tcf7l2*-cKO mice, and with both genotypes showing a marked day-to-day reduction in chasing events, indicative of social habituation (Fig. 5I, violin plot; Fig. 5J, heatmap; SFig. 10D, bar plot; compare with Fig. 3H-I and SFig. 5E). The diet, however, had no effect on grouping preferences: while control mice exhibited more flexible spatial use during social interactions, *Tcf7l2*-cKO mice retained rigid preferences for specific social grouping within defined locations (Fig. 5K-L).

In a group of unfamiliar males (Fig. 5M), the ketogenic diet reduced the number of spontaneous social investigative interactions by approximately 30-40% (Fig. 5N; compare with Fig. 3M), resembling the reduction in social exploration observed in females tested in the *EcoHAB* system. The short-time-dependent decline in investigative interactions observed under the standard diet was abolished after ketogenic diet treatment, with both *Tcf7l2*-cKO and control males maintaining a relatively stable number of interactions across time bins.

In conclusion, the ketogenic diet effectively normalized energy metabolism in the thalamocortical circuitry of TCF7L2-deficient mice. While it influenced behavior in control animals, it also alleviated certain behavioral deficits in *Tcf7l2*-cKO mice, supporting a possible link between brain metabolic dysfunctions and impairments in social behavior and locomotor patterns.

## Discussion

We uncovered that TCF7L2 contributes to social behavior and locomotor flexibility via its thalamic functions. We also demonstrated that TCF7L2 regulates energy metabolism in the thalamocortical circuit by modulating metabolic gene expression in the thalamus. Furthermore, we showed that dietary intervention can alleviate both brain metabolic impairments and some behavioral alterations caused by TCF7L2 deficiency.

### Shared risk gene mutations as a cause of comorbidity between metabolic and psychiatric disorders

Our results highlight that the frequent comorbidity between systemic metabolic and psychiatric disorders may stem from mutations in shared risk genes, with *TCF7L2* representing one such factor. Misregulation of *Tcf7l2* in metabolic organs leads to glucose intolerance and insulin resistance, while disrupting *Tcf7l2* in postnatal thalamic neurons alters energy metabolism in the thalamocortical circuitry and leads to social deficits. Notably, this conclusion is consistent with a recent human study that identified TCF7L2 as a key regulator in protein networks dysregulated in metabolic and mental disorders ^80^.

While numerous models exist for either metabolic impairments or mental disorders, integrated models that reflect the co-occurrence of both conditions are rare. Our *Tcf7l2* knockout model combines postnatal impairments in the thalamocortical system with poor glucose control in the periphery likely due to a partial knockout in the pancreas alongside an increase in TCF7L2 levels in the liver. This model aligns with existing models of TCF7L2-associated metabolic impairments in the periphery because high fasting plasma glucose levels or glucose intolerance have been observed in mice with either pancreatic knockout of *Tcf7l2* or *Tcf7l2* overexpression in the liver ^19,81–84^. Consequently, our study offers a useful model for exploring the interaction between systemic metabolic disorders and mental disorders associated with thalamocortical dysfunction.

### Thalamocortical impairment as a cause of autism spectrum disorder

Symptoms frequently accompanying ASD—sensory processing abnormalities, impaired attention, seizures, and disrupted sleep—have been observed in mouse models carrying mutations in ASD risk genes (*Shank3*, *Tsc1*, *Ptchd1*, and *Cntnap2*), in which thalamic dysfunction has been implicated ^59–62^. A recent study focusing on the thalamic reticular nucleus, an inhibitory structure that regulates the activity of the proper thalamus, reported social deficits ^63^. These findings indicate that thalamocortical dysfunction may primarily contribute to ASD-related phenotypes. Our results extend this framework by demonstrating that reduced social interest—a core ASD-related feature—as well as locomotor inflexibility can also originate from thalamic circuit disruptions, even when these occur postnatally, as in our model. This highlighting the broader role of the thalamus and its developmental trajectory in ASD pathophysiology.

Our findings revealed a complex pattern of behavioral alterations in *Tcf7l2*-cKO mice. On one hand, these animals exhibited diminished interest in social stimulus, reduced social chasing behavior (a key component of hierarchy formation ^68^), rigid social grouping preferences and a failure to respond to social cues. Collectively, these deficits point to impaired group-level social dynamics. On the other hand, *Tcf7l2*-cKO mice showed decreased general exploration and impairments in locomotor habituation and adaptation. These observations suggest that postnatal loss of TCF7L2 in the thalamus leads to reduced behavioral flexibility in response to changing environment and social contexts.

These findings raise the question of which specific thalamic nuclei may underlie the observed behavioral deficits. Identifying the specific thalamic nuclei responsible for social behaviors remains challenging. Higher-order nuclei may be involved, such as those that interact with the prefrontal cortex or nonspecific nuclei project to multiple brain regions. Indeed, the nucleus reuniens, posterior intralaminar complex and the mediodorsal nucleus have been linked to social behaviors in mice ^52–58^. Notably, the reduced engagement in chasing behavior in our model may point to involvement of the mediodorsal thalamus, as activity within the mediodorsal thalamus–prefrontal cortex circuit has been shown to play a critical role in regulating social dominance ^52,53^. Another possibility is that these deficits result from generalized sensory filtering issues within the thalamus. Sensory dysregulation is known to emerge early in the progression of ASD and is strongly implicated in social functioning deficits ^85,86^. Further studies are needed to clarify which thalamic regions contribute to TCF7L2-related social impairments and to uncover the underlying mechanisms.

### Linking energy metabolism to the etiology of mental disorders

Our findings adds to the growing body of evidence that dysregulated energy metabolism in the brain plays a crucial role in the etiology of mental disorders ^87,88^. In particular, our data point to a relationship between altered glucose metabolism within thalamocortical circuits and ASD-relevant behavioral phenotypes, although a causal link remains to be established. This aligns with clinical studies showing elevated levels of glycolysis products—pyruvate, alanine, and lactate—in the blood of individuals with ASD, as well as elevated cerebral lactate levels ^89–94^, suggesting enhanced aerobic glycolysis. Similarly, our model exhibited elevated lactate and alanine levels alongside reduced pyruvate oxidation in the thalamus and cortex. This indicates a shift in energy metabolism from glucose oxidation toward alternative energy substrates such as fatty acids and ketone bodies. The *Tcf7l2*-cKO knockout model thus provides experimental support for the Warburg effect hypothesis in ASD ^94^ and establishes these mice as a relevant model for studying metabolic dysfunctions in ASD and other mental disorders.

At the molecular level, these metabolic alterations observed in the thalamocortical circuit in our mouse model may reflect dysregulation of gene networks involved in lipid and glucose metabolism. In particular, thalamic TCF7L2 deficiency altered the expression of genes involved in lipid metabolism, including the key transcriptional regulators PPARD and RORA, as well as glucose metabolism regulators such as PKCδ; the latter two are enriched in the thalamus (see mouse.brain-map.org). The PKCδ is a likely regulator of the pyruvate dehydrogenase complex ^73,74^, which catalyzes the critical step for pyruvate entry into the TCA cycle. In *Tcf7l2* knockout mice, we observed reduced levels of PKCδ alongside increased phosphorylation of PDHEα, providing preliminary evidence for the disruption of the hypothetical TCF7L2-PKCδ-PDHc regulatory axis.

Strikingly, this metabolic disruption affected the thalamus and the cortex, even though the knockout was restricted to the thalamus. This may be attributed to the extensive synaptic connections between thalamic neurons and cortical regions. Another possibility is that brain effects are a consequence of changes in systemic metabolism. However, two observations argue against this possibility. First, the cerebellum, which lacks direct thalamic input, showed no metabolic abnormalities, pointing to the specificity of changes in the thalamocortical circuit. Second, while the ketogenic diet normalized metabolite oxidation in the thalamus and cortex, it did not improve peripheral glucose homeostasis, suggesting that the observed brain effects were not driven by systemic metabolic alterations.

These circuit-level metabolic alterations may have direct clinical relevance. Reduced activity of the PDH complex in the thalamocortical circuit may represent a convergent point in the etiology of a subset of individuals with ASD and SCZ, given that the gene encoding the key PDHE1α (PDHA1) subunit of the PDHc is an ASD-risk gene, and autoantibodies against this protein—potentially disrupting PDHc activity—are found in some patients with SCZ ^95^. This disruption in glucose metabolism offers a potential target for therapy, as the ketogenic diet, already used in individuals with congenital pyruvate dehydrogenase deficiency ^96,97^, has shown effectiveness in *Tcf7l2*-cKO mice, supporting its therapeutic potential for a specific group of patients. However, it is important to recognize that, despite its expected benefits for brain metabolism demonstrated in the present study, long-term use of the ketogenic diet might have side effects in both mouse models ^98,99^ and humans ^100^.

In our model, we observed a co-occurrence of metabolic impairments in the brain and behavioral abnormalities. However, electrophysiological disturbances have also been identified in the same model ^31^, and other alterations may be present as well. Thus, it remains unclear whether the metabolic deficits are causally related to the observed behavioral phenotypes. The fact that the ketogenic diet led to improved brain metabolism and behavior provides a potential clue, although the nature of this relationship remains to be established. Investigating the relationship between *Tcf7l2*-related metabolic alterations in the brain and behavioral deficits will be the focus of future studies.

## Conclusion

In conclusion, our findings highlight a potential link between disrupted energy metabolism in the thalamocortical circuitry and the etiology of mental disorders such as ASD and SCZ, while emphasizing the role of TCF7L2 in these processes and expanding its known role beyond regulating peripheral metabolism. The observed metabolic shift in energy substrate utilization, with reduced pyruvate oxidation due to thalamic TCF7L2 deficiency, suggests that metabolic dysregulation may directly contribute to neuropsychiatric symptoms. Furthermore, our findings demonstrate that a ketogenic diet can restore both metabolic and social deficits in *Tcf7l2* knockout mice, providing preliminary evidence that dietary interventions may help mitigate the metabolic contributions to mental disorders.

## Materials and Methods

### Animals

Thalamus-specific *Tcf7l2* conditional knockout (cKO) mice (*Cck^Cre^:Tcf7l2^fl/fl^*) were generated as previously described ^31^. Briefly, the C57BL/6NTac-*Tcf7l2*^tm1a(EUCOMM)Wtsi^ (*Tcf7l2*^tm1a^) mouse strain, which contained with a trap cassette upstream of the critical exon 6 of the *Tcf7l2* gene, was first crossed with flippase-expressing mice (Gt(ROSA)26Sor^tm1(FLP1)Dym^; Jackson Laboratory, #009086), and then with mice which expressing Cre recombinase under the control of the *Cck* promoter (*Cck*^tm1.1(cre)Zjh/J^; Jackson Laboratory, #012706). *Cck^Cre:^Tcf7l2^+/+^* animals were used as controls. To generate the *Cck^Cre^:TdTomato^fl/+^* reporter strain, *Cck^Cre^:Tcf7l2^+/+^* mice were crossed with the Ai9(RCL-tdT) strain (Gt(ROSA)26Sor^tm9(CAG-tdTomato)Hze/J^ Jackson Laboratory, #007909). All animals were housed under standard laboratory conditions (21 ± 2 °C, 60% ± 10% humidity, and 12 h/12 h light/dark cycle), with food and water provided *ad libitum*. Mice aged P60–P90 were used for the tests, unless stated otherwise. All experimental procedures complied with the normative standards of the European Community (Directive 86/609/EEC) and the Polish Government (Dz.U. 2015 poz. 266). Protocols were approved by the Local Ethical Committee No. 1 in Warsaw, and the institutional animal welfare advisory board oversaw animal use at the Centre of New Technologies.

### Western blot analysis

Organs and selected brain structures (n = 4 per group) were dissected and homogenized in ice-cold RIPA buffer (50 mM Tris, pH 7.5, Bioshop Life Science Products, #TRS001.1; 50 mM NaCl, Chempur, #794121116; 1% NP-40, Thermo Scientific, #85124; 0.5% sodium deoxycholate, Merck Life Science, #D6750; 0.1% sodium dodecyl sulfate [SDS], Bioshop Life Science Products, #SDS999.500; 1 mM ethylenediaminetetraacetic acid [EDTA], VWR, #E177; 1 mM NaF, Merck Life Science, #67414-1ML-F; EDTA-free protease inhibitor cocktail cOmplete, Roche, #4693132001; phosphatase inhibitor PhosSTOP^TM^, Roche, #4906845001). Protein concentrations were measured using the Protein Assay Dye Reagent (Bio-Rad Laboratories, #5000006). Clarified protein homogenates were separated on SDS-polyacrylamide gels (Bio-Rad, #1610183) and transferred to nitrocellulose membranes (Bio-Rad, #1620112). Membranes were blocked with 10% nonfat dry milk in phosphate-buffered saline (PBS) containing 0.1% Tween 20 (PBS-T) (VWR, #663684B), and incubated overnight at 4 °C with the following primary antibodies: anti-TCF7L2 (Cell Signaling Technology, #2569, 1:1000), anti-GAPDH (Santa Cruz Biotechnology, #sc-25778, 1:500), anti-PKCδ (Thermo Fisher Scientific, #PA5-17552, 1:1000), anti-PDHE1α (Santa Cruz Biotechnology, #sc-377092, 1:30000), anti-phospho-Ser232-PDHE1α (Merck KGaA, #AP1063, 1:500). After washing, membranes were incubated for 2 h at room temperature with secondary antibodies: anti-rabbit IgG peroxidase (Sigma Aldrich, #A0545, 1:10000) or anti-mouse IgG peroxidase (Sigma Aldrich, #A9044, 1:10000). Signal detection was performed using chemiluminescence, and images were captured with an ImageQuant LAS 4000 (Cytiva). Densitometric analyses were conducted using Quantity One 1-D software (Bio-Rad).

### Tissue fixation

Mice were anesthetized with a mixture of ketamine (Biowet, 125 mg/kg body weight) and xylazine (Biowet, 10 mg/kg body weight). They were then transcardially perfused with 0.1 M PBS, followed by 4.5% paraformaldehyde (PFA; Merck Life Science, #P6148) in PBS. The brain, liver, duodenum, and pancreas were dissected, incubated overnight in 4.5% PFA, and saturated in 30% sucrose (Merck Life Science, #1076515000) in PBS at 4 °C for 24 h. The tissues were subsequently transferred into O.C.T. compound (Sakura Tissue-Tek, #4583) and frozen in-30 °C isopentane (VWR, #24872.298). Organ samples were sectioned using a Leica CM1860 cryostat. Brain sections (30 μm thick) were collected in an anti-freeze solution containing 30% sucrose (Merck Life Science, #1076515000) and 30% glycerol (VWR, #443320113) in 0.1 M PBS, pH 7.4. Liver, duodenum, and pancreas sections (15 μm thick) were mounted on glass slides.

### Immunohistochemistry

Frozen sections (n = 4 per group) were washed in 0.2% Triton X-100 in PBS (PBS-T), incubated in 0.3% hydrogen peroxide (H2O2) for 10 min, blocked with 5% normal goat serum in PBS-T, and then incubated overnight at 4 °C with anti-TCF7L2 antibody (Cell Signaling Technology, #2569, 1:1000) in 1% NGS. Next, sections were incubated for 1 h with biotinylated goat anti-rabbit antibody (Vector Laboratories, #BA-1000, 1:500) in 1% NGS, and then for 1 h in Vectastain ABC reagent (Vector Laboratories, #PK-6100). Staining was developed using 0.05% DAB (Sigma-Aldrich, #D12384) and 0.01% H2O2. Finally, sections were mounted onto SuperFrost Plus slides, dehydrated in an ethanol series (50%, 70%, 95%, and 99.8%), washed in xylene, and mounted using EuKitt.

### Immunofluorescence

Frozen sections (n = 4 per group) were washed in PBS-T and blocked with 5% normal donkey serum (NDS) for 1 h, and then incubated with anti-TCF7L2 antibody (1:500; 2569, Cell Signaling Technology) in 1% NDS overnight at 4 °C. Sections were then incubated for 1 h with a secondary antibody conjugated with Alexa Fluor 488 (donkey anti-rabbit immunoglobulin G [IgG] with Alexa Fluor 488, Invitrogen, #A-21206). Slides were additionally stained with Hoechst 33342 (1:10,000; #62249, Thermo Fisher Scientific), washed, and mounted with Vectashield Antifade Mounting Medium (H1000, Vector Laboratories). Images were captured with an Axio Imager Z2 LSM 700 Zeiss confocal microscope.

### Diet intervention

Adult young mice (postnatal day 40-50) were fed a standard chow diet (Ssniff® - E1500-04; energy: 11% fat, 23% proteins, 66% carbohydrates), high-fat diet (Ssniff® - E15742-34; energy: 60% fat, 20% proteins, 20% carbohydrates) or ketogenic diet (Ssniff® E15149-30; energy: 94.3% fat, 4.5% proteins and 1.2% carbohydrates) for 6 weeks.

### General metabolic assessment: body weight, glucose, and insulin blood level, intraperitoneal glucose tolerance test

Mice weight was measured at the beginning and end of the diet intervention (n = 6 per group). Plasma was collected for insulin and glucose measurements. Plasma insulin (n = 10 controls and 15 *Tcf7l2*-cKO mice per group) was measured using the insulin ELISA kit (Merck, Sigma-Aldrich #EZRMI-13K), and glucose (n = 8 controls and 9 *Tcf7l2*-cKO mice per group) using the glucose oxidase method (BioSystems, #21503) in animals fasted overnight. Glucose tolerance (n= 6 per group) was assessed by intraperitoneal glucose injections (2g/kg body weight) to animals fasted overnight. Blood samples were collected from the tail at 0, 15, 30, 60, 90, and 120 min after glucose injection.

### Basic locomotor activity

Locomotor activity was assessed in a dimly lit three-chamber apparatus enriched with objects during a 10-min session (n = 9 per group). Movement parameters were automatically quantified using AnyMaze behavioral tracking software (Stoelting, Wood Dale, IL).

### Light-dark box

A light/dark transition test (n = 9 per group) was conducted using a cage divided into two equal compartments, with a small opening allowing free movement between them (Ugo Basile, Italy). The open (light) compartment faced the wall to prevent the animals from seeing the experimenter. Illumination in the open compartment was set to 200-400 lux, while the dark compartment was maintained at ≤5 lux. Each animal was placed in the center of the light compartment and allowed to explore the cage freely for 10 min. After each trial, urine and fecal boli were removed, and the cage was cleaned with 70% ethanol to eliminate potential olfactory cues. Recordings were analyzed to extract latency to enter the dark compartment, time spent in the light and dark compartment, and the number of transitions between them.

### Open-field test

An open-field test (n = 10 females per group and 8 males per group) was performed in a dark room using a black-painted square arena (90 x 90 cm), enclosed by walls (30 cm high). The center was illuminated with a 50 W halogen bulb suspended centrally 30 cm above the arena. The animal was placed in the thigmotaxic zone, facing one of the corners, and its movement was tracked for 10 min. The recordings were then analyzed to extract data on time, distance moved, and movement duration in the illuminated central zone (a circle directly corresponding to the brightly lit part of the arena) and the thigmotaxic zone (perimeter).

### Buried food test

The buried food (n = 8 per group) test was used to test the olfactory function in mice. Before the test, a piece of Kellogg’s Fruit Loops was added to the subjects’ cages for three days. All chow pellets were removed 18-24 hours before the test. The test was conducted in a standard mouse cage with a 3 cm layer of fresh bedding. On the test day, each mouse was acclimated to the test cage for 5 min, then transferred to an empty, clean cage. The food stimulus was buried approximately 1 cm beneath the surface of the bedding in a random corner of the test cage. The mouse was then returned to the test cage, and the latency to locate the buried food was recorded.

### Three-chamber test

The three-chamber social preference test (n = 9 per group) was conducted according to previous recommendations ^101^. The test consisted of three distinct phases of 10 min each. The first was the habituation phase, during which the test mouse could explore the apparatus containing two empty cups. The second stage was the pre-test, where two identical objects were placed under the cups to familiarize the animal with the objects and to observe any possible preference for a specific chamber. Finally, the social preference phase allowed the test mouse to interact with a social stimuli - an age-, sex-, and strain-matched unfamiliar WT mouse under one cup - and a non-social stimulus - a new object under the other cup. The test mouse was recorded for 10 min, and the time spent interacting with the social and the non-social stimulus was quantified. The area directly surrounding each cup was designated as a zone of interest, and the mice’s presence in this area was automatically scored for social preference using the Anymaze behavior tracking software (Stoelting, Wood Dale, IL). All videos were manually reviewed to verify that the animal was tracked well, and the time when the animal was near the cup without any interaction with the mouse or the object was removed from the final analysis. Social preference was quantified as the discrimination index: (Time exploring the mouse - Time exploring the object) / (Total time spent exploring).

### Electronic tagging and pre-experimental housing for IntelliCage and EcoHAB

Female mice were electronically tagged under brief isoflurane anesthesia by subcutaneous injection of glass-covered microtransponders (11.5 mm length, 2.2 mm diameter, DATAMARS for *IntelliCage*, or 9.5 mm length, 2.2 mm diameter, RFIP Ltd for *EcoHAB*) for individual identification and tracking. The microtransponders emit a unique animal identification code when activated by the magnetic field of RFID antennas or a portable chip reader. After tagging, mice were transferred to the experimental rooms and housed together for at least one week prior to placement in the *IntelliCage* or *EcoHAB* systems, in order to minimize the effects of an unstable social structure and to acclimate to the shifted light/dark cycle. The temperature in the experimental rooms was maintained at 23–24 °C, with relative humidity levels between 35% and 45%.

### IntelliCage

The automated *IntelliCage* system consisted of a home cage equipped with four conditioning chambers (corners), each fitted with two water bottles, sensors, and motorized door. The cognitive assessment in this setup involved two independent cohorts per genotype, n = 23 for control and n = 18 for *Tcf7l2*-cKO female mice. The experiment began with a three-day simple adaptation phase (SA), followed by a two-day nose-poke adaptation phase (NP), a three-day place learning phase (PL), and a reward-based discrimination learning phase (DISC). During SA, doors in all conditioning units (corners) were open, and access to the two bottles in each corner, filled with tap water, was unrestricted. During NP, the doors were closed by default, and the mice were required to use a nose-poke response—by inserting their snouts into any of the two holes on the operant corner walls to gain access to water. The door remained open as long as the animal kept its snout in the hole, regardless of drinking behavior. In PL, each mouse was randomly assigned to a conditioning unit that provided access to water. Only one unit was available per mouse, and no more than three mice were assigned to the same unit. In DISC, one bottle in each corner was filled with 10% sucrose. Mice could choose between nose-poking one of two separate doors—one leading to a bottle of tap water and the other to a bottle containing a reward—placed in opposite locations within the conditioning corner. All visits and nose-pokes were recorded. Nose-pokes directed to the bottle with reward as the first choice upon entering the conditioning unit were considered a correct responses. Animals exhibiting transponder identification failure were excluded from the analysis.

### Eco-HAB

This naturalistic automated system *EcoHAB* ^67^ consisted of four cages connected by tunnels, arranged in a looped square layout, with food and water available in two of them. The behavioral assessment in this setup involved cohorts of control and *Tcf7l2*-cKO female mice (three independent cohorts fed the chow diet, n = 35 for control and n = 32 for *Tcf7l2*-cKO mice; two independent cohorts fed the ketogenic diet, n = 21 for control and n = 20 for *Tcf7l2*-cKO mice). All mice in a tested cohort had unrestricted access to all compartments throughout the 3-day experimental period. Individual mouse movements between cages were tracked using RFID antennas installed on the connecting tunnels. Daily movement data were processed using the custom ‘DeepEcoHab’ software package (see Code and data availability section) to extract relevant behavioral features. Exploratory behavior was quantified as the total number of visits each mouse made to all EcoHAB cages. Dynamic social interactions were assessed by measuring chasing behavior between pairs of mice. In-pair chasing was defined as an event where one mouse entered a tunnel, followed immediately by a second mouse entering the same tunnel and both traveling together in the same direction until exiting into the next cage. Passive social interactions were assessed as time spent together, defined as the total duration during which both mice occupied the same cage, calculated for each pair. To assess grouping behavior, this total time was then expressed as the percentage of time the pair spent together in each individual cage. Animals exhibiting transponder identification failure during the experiment were excluded from the analysis.

### Social investigative interactions between unfamiliar mice

The spontaneous social interaction test was conducted in males. Four age-and strain-matched unfamiliar mice were placed together into a test cage (*n* = 16 per group in 4 independent cohorts), and their behavior was recorded continuously for 2 hours. Nose-to-nose contact and anogenital sniffing ^102^ were scored and quantified from the recordings.

### Metabolic substances extraction and 1D NMR measurements

Soluble metabolites were extracted from the thalamus and cortex (n = 6 per group) and separated using Bligh and Dyer method ^103^. Samples were homogenized on ice with 1 ml chloroform:methanol (2:1, v/v) containing 0.01% butylated hydroxytoluene as an antioxidant. After adding 0.5 ml water and mixing, samples were centrifuged (3000 rpm, 10 min, 4 °C) to separate phases. The aqueous phase was frozen in liquid nitrogen and freeze-dried, then dissolved in deuterium oxide with 0.05% 2,2-dimethyl-2-silapentane-5-sulfonate (DSS) as an internal standard and transferred into 3 mm NMR tubes. All one-dimensional ^1^H-NMR measurements were performed on the 700 MHz Agilent DirectDrive2 spectrometer equipped with a room-temperature HCN probe, temperature-controlled at 25 °C. ^1^H-NMR spectra were recorded with 32K data points, acquisition time 3s, interscan delay 90s and 1250 scans. The obtained 1D data sets were processed and analyzed using Mnova 12.0 software with zero-filling to 128K and exponential weighting function (line broadening of 0.2 Hz). Zero-order phase correction and baseline correction using the polynomial fit were done automatically on each spectrum. The chemical shift scale was referenced to the 4,4-dimethyl-4-silapentane-1-sulfonic acid (DSS) peak (δ = 0.00 ppm). Spectra were used for metabolite identification. Lactate (Lac), alanine (Ala), glycine (Gly), asparagine (Asp), succinate (Suc), γ-aminobutyric acid (GABA), taurine (Tau), N-acetylaspartic acid (NAA), creatine (Cr), myo-inositol (M-In) and malic acid (MA) were identified ^104,105^. Absolute integrals of representative peaks were determined.

### High-resolution respirometry

Oxygraph-2k high-resolution respirometer (Oroboros Instruments, Innsbruck, Austria) was used to measure mitochondrial oxidation by substrate-driven respiration protocol ^106^ in samples from the thalamus, cortex, and cerebellum (n = 3 per group). Tissues were permeabilized with 0.005% digitonin for 30 minutes before the start of the respirometry assy. Substrates were added sequentially in the following order: malate (0.5 mM), palmitoylcarnitine (0.5 μM), ADP+Mg^+2^ (2 mM), octanoylcarnitine (0.2 mM), 3-hydroxybutyrate (1 mM), pyruvate (5 mM), glutamate (10 mM), and succinate (10mM). The integrity of the outer mitochondrial membrane was verified by the addition of cytochrome c (10 µM) after the addition of all substrates. Maximum respiration was assessed through stepwise addition of carbonyl cyanide m-chlorophenyl hydrazone (CCCP 0.05 µM). Complex I was blocked by rotenone (0.5 µM). Complex II was inhibited by malonate (5 mM). Complex III was inhibited by Antimycin A (2.5 µM). Complex IV was inhibited by sodium azide (100 mM). MIR-06 buffer was used to access respiration (ethylene glycol-bis(β-aminoethyl ether)-N,N,Nʹ,Nʹtetraacetic acid [EGTA] 0.5 mM, MgCl2 3 mM, lactobionic acid 60 mM, taurine 20 mM, KH2PO4 10 mM, HEPES 20 mM, d-sucrose 110 mM, both 1 mg/ml BSA–fatty acid free and Catalase 280 U/mL were added freshly, pH 7.1). Palmitoylcarnitine was prepared in the presence of BSA, and preliminary experiments were conducted to ensure the applied concentration did not inhibit respiration. Due to limited sample availability, all substrates were added within a single experimental run. Data was recorded and analyzed with DatLab software (v5.1.0.20; Oroboros Instruments, Innsbruck, Austria).

### Statistical analysis

All experiments were randomized and analyzed in a blinded manner. The required sample sizes were estimated based on our previous experience with similar experiments and articles published in the same research field. The indicated *n* for each experiment represents the number of biological replicates. Data were tested for normality (Shapiro-Wilk) and further analyzed using GraphPad Prism 9.5.1. Statistical tests were chosen based on the experimental design and data distribution. Two-tailed Mann-Whitney test was used to compare two independent groups when data did not meet the assumptions of parametric tests. Two-tailed T-test was used to compare the means of two independent groups when data were normally distributed and variances were equal. One-way analysis of variance (ANOVA) was used to compare the means of three or more independent groups. Two-way ANOVA was used to analyze the effects of two independent variables on a dependent variable. Repeated measures ANOVA was used for time-series data. Correlations between parameters were assessed using Pearson’s correlation coefficient. Differences in variance between groups were evaluated with Levene’s test. In all cases, statistical significance was considered when *p* < 0.05.

### Code and data availability

The algorithms and scripts comprising the’DeepEcoHAB’ package - the software implementation used to generate and analyze *EcoHAB* results reported herein - are currently under final development and are not yet publicly accessible. The package will be released under an Open Source Initiative-approved open-source license, permitting free academic use via GitHub (URL to be provided) in the last quarter of 2025. Until that release, the code is available from the coauthors of the study: Marcin Lipiec & Konrad Danielewski, upon reasonable request. Raw data from *EcoHAB* test will be made available in the open-access data repository of the University of Warsaw.

## Acknowledgements

This work was supported by Narodowe Centrum Nauki (NCN, National Science Center) grants 2020/37/B/NZ4/03261 and 2024/55/B/NZ4/03078 to MBW and 2015/19/B/NZ4/03571 to AN. Additionally, ACh and BHMM were supported by NCN grant 2019/35/B/NZ3/04066 awarded to ACh.

