## Supplemental Material for "The Role of TCF7L2 in Regulating Energy Metabolism in Thalamocortical Circuitry and its Broader Impact on Social Behavior"

#### Supplementary Figure 1

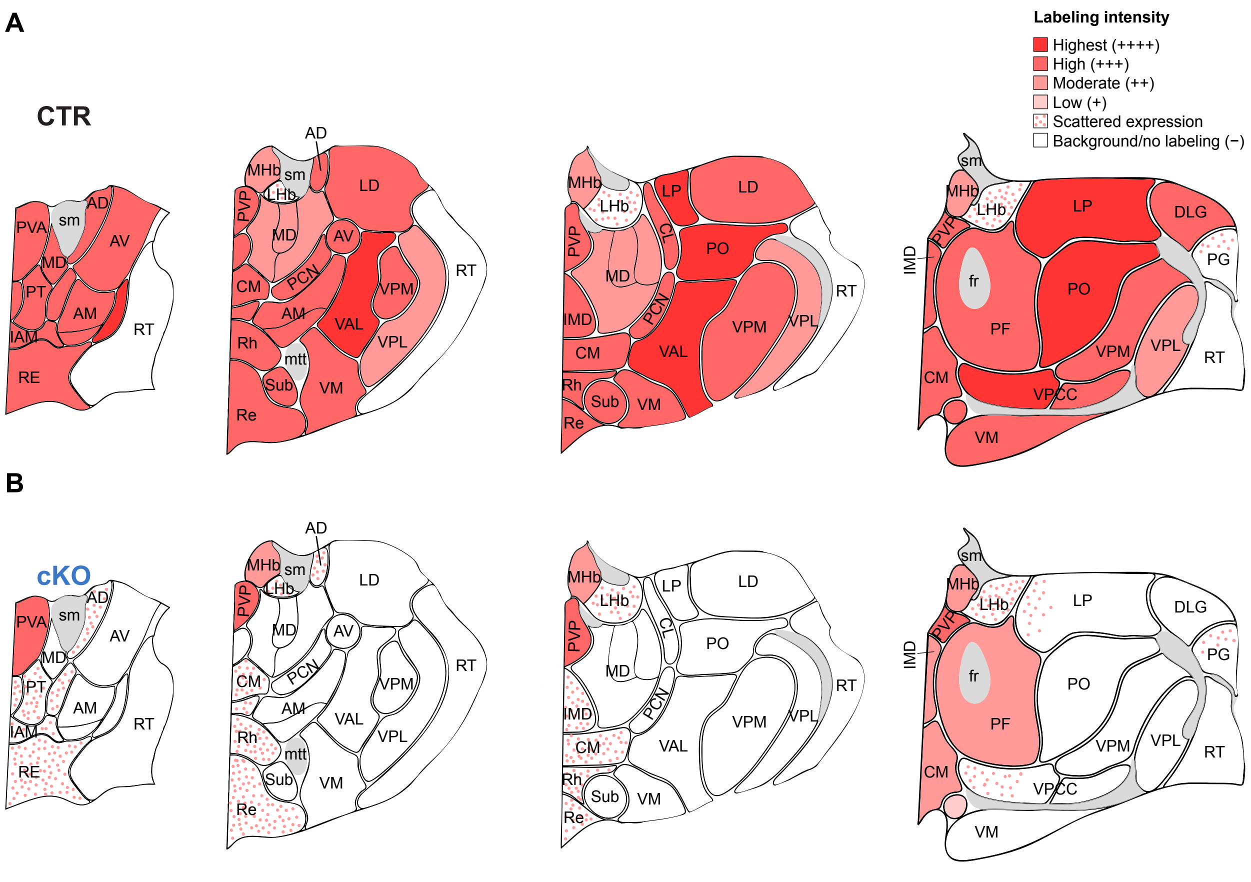


**SFig. 1. TCF7L2 is expressed in thalamic nuclei, and its expression is lost in mice with postnatal thalamic *Tcf7l2* knockout**

**(A-B)** Schematic representation of DAB immunohistochemical staining of TCF7L2 in coronal sections of the adult brain from (A) control (CTR; based on Nagalski et al., 2016 [29]) and (B) *Cck*$^{Cre}$*:Tcf7l2*$^{fl/fl}$ (cKO; based on Lipiec et al., 2020 [31]) mice.

AD, anterodorsal thalamic nucleus; AM, Anteromedial thalamic nucleus; AV, Anteroventral thalamic nucleus; CL, centrolateral thalamic nucleus; CM, central medial thalamic nucleus; DLG, dorsal lateral geniculate nucleus; fr, fasciculus retroflexus; IMD, intermediodorsal thalamic nucleus; LD, laterodorsal thalamic nucleus; LHb, lateral habenula; LP, lateral posterior thalamic nucleus; MD, mediodorsal thalamic nucleus; MHb, medial habenula; PF, parafascicular thalamic nucleus; PG, pregeniculate nucleus; Po, posterior thalamic nuclear group; PVA, paraventricular thalamic nucleus, anterior part; PVP, paraventricular thalamic nucleus, posterior part; Re, reuniens thalamic nucleus; Rh, rhomboid thalamic nucleus; RT, reticular thalamic nucleus; Sub, submedius thalamic nucleus; sm, stria medullaris; VA/VL, ventral anterior/ventral lateral thalamic nuclei; VM, ventromedial thalamic nucleus; VPM, ventral posteromedial thalamic nucleus; VPL, ventral posterolateral thalamic nucleus; VPPC, ventral posterior nucleus of the thalamus, parvicellular part.

#### Supplementary Figure 2

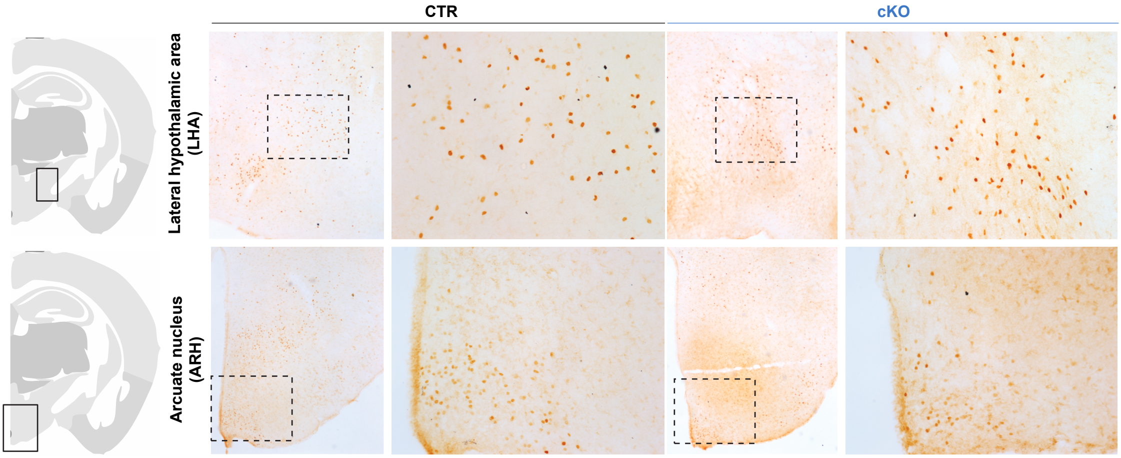


**SFig. 2. TCF7L2 depletion in the thalamus does not affect hypothalamic TCF7L2 expression**

DAB immunohistochemical staining of TCF7L2 in coronal brain sections from control (CTR) and *Cck*$^{Cre}$:*Tcf7l2*$^{fl/fl}$ (cKO) male mice showing the hypothalamic area.

#### Supplementary Figure 3

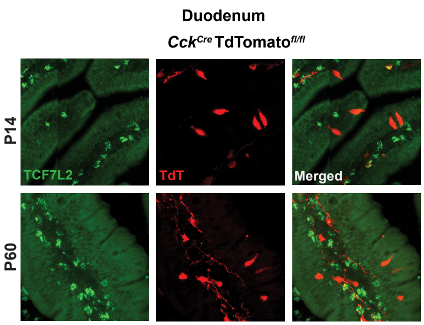


**SFig. 3. *Cck*-driven Cre activity does not target TCF7L2-expressing cells in the duodenum**

Immunofluorescent staining of TCF7L2 (green) and TdTomato (red) in the duodenum of the $Cck^{Cre}$*:Td-Tomato*$^{fl/fl}$ reporter line at P14 and P60, illustrating that Cre-driven TdTomato expression does not colocalize with TCF7L2-positive cells.

### Supplementary Figure 4

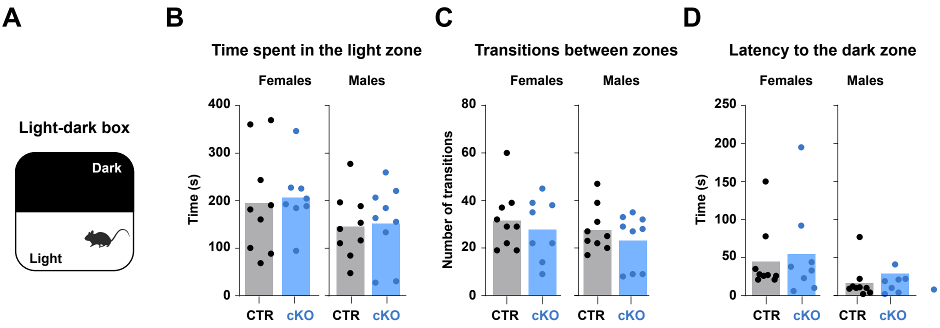


**SFig. 4. TCF7L2 depletion in the thalamus does not affect anxiety levels**

**(A)** Schematic of the light-dark box test; both sexes were tested. **(B)** Total time spent in the light zone by control (CTR) and *Cck*$^{Cre}$*:Tcf7l2*$^{fl/fl}$ (cKO) mice. **(C)** Number of transitions between zones. **(D)** Latency to enter the dark zone for female and male mice.

Bars in graphs represent mean values, with dots indicating individual mice. Data in B, C, and D were analyzed with two-way ANOVA, followed by Tukey's multiple comparison test.

### Supplementary Figure 5

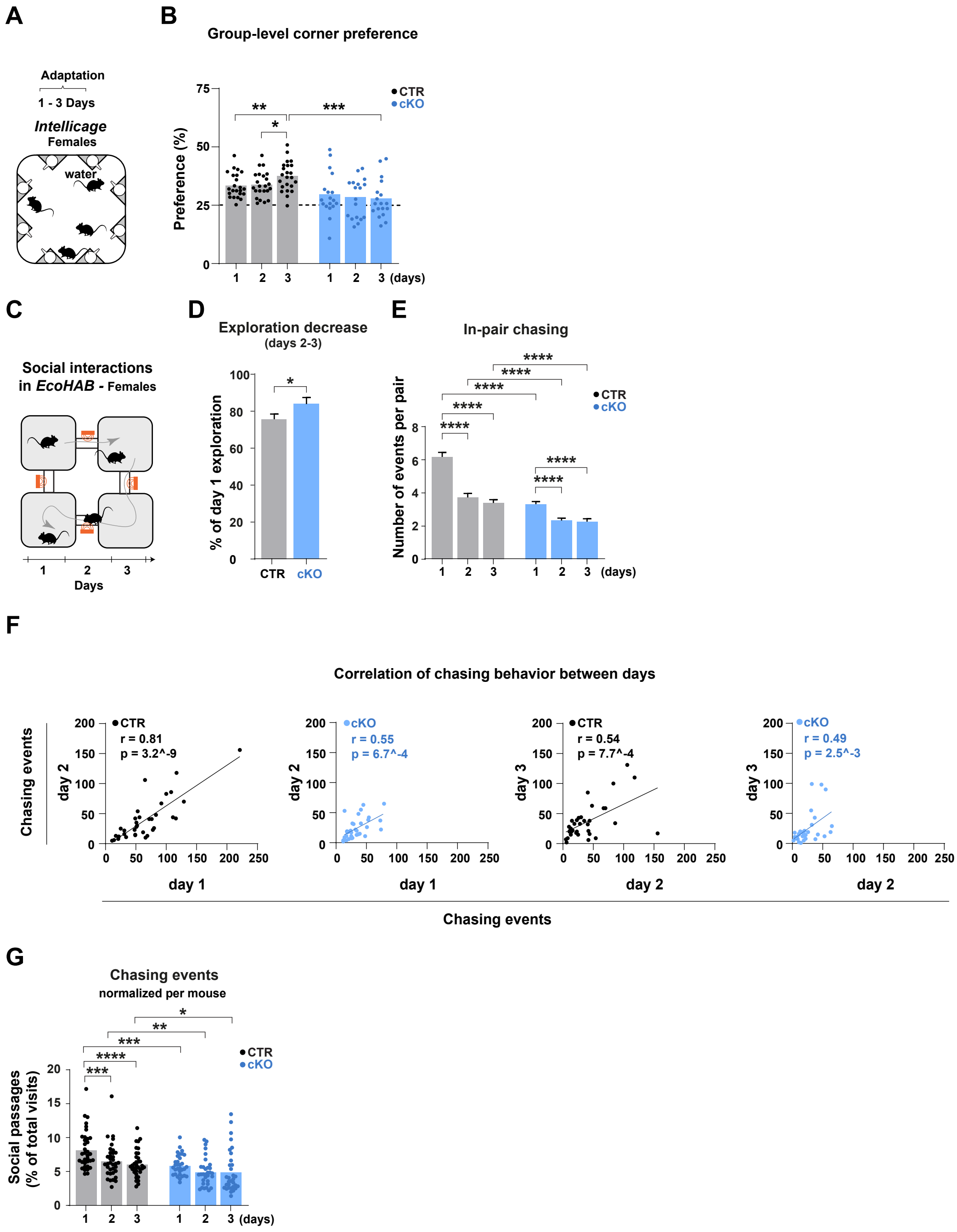

**SFig. 5. TCF7L2 depletion in the thalamus disrupts coordinated group behaviors**

**(A)** Schematic of the *Intellicage* setup showing water access in four corners during the simple adaptation phase (days 1-3). **(B)** Percentage of each mouse's visits to the group's most preferred corner, relative to its total number of visits, shown for control (CTR) and $Cck^{Cre}$:$Tcf7l2^{fl/fl}$ (cKO) mice; CTR n = 23 and cKO n = 18 in two independent cohorts for each genotype; only females were tested; results irepresent data from the dark phase. **(C)** Schematic of the EcoHAB setup used to assess social behavior. CTR n = 35 and cKO n = 32 in three independent cohorts for each genotype; only females were tested; results in D-G represent data from the dark phase. **(D)** Percentage decrease in exploratory activity (visits to all cages) in days 2-3 relative to day 1. **(E)** In-pair chasing events, measured as one mouse trailing another through a tunnel for each mouse pair; the bar plot corresponds to the violin plot shown in Fig. 3H. **(F)** Correlation analyses of chasing events across consecutive days (days 1–2 and 2–3) for control (CTR) and $Cck^{Cre}$:$Tcf7l2^{fl/fl}$ (cKO) mice. **(G)** Percentage of social passages (chasing and being chased) relative to total cage visits.

Bars in B, D, E and G represent mean values. Dots in B, F and G indicate individual mice. Error bars in D and E represent standard deviation. Data in B, E, and G were analyzed using repeated measures two-way ANOVA, followed by Tukey's multiple comparison test; data in D were analyzed using an unpaired T-test. In B, D, E and G, *$p < 0.05$, **$p < 0.01$, ***$p < 0.001$, ****$p < 0.0001$. Data in F were analyzed using Pearson´s correlation test; r- and p-values are shown on the plots.

### Supplementary Figure 6

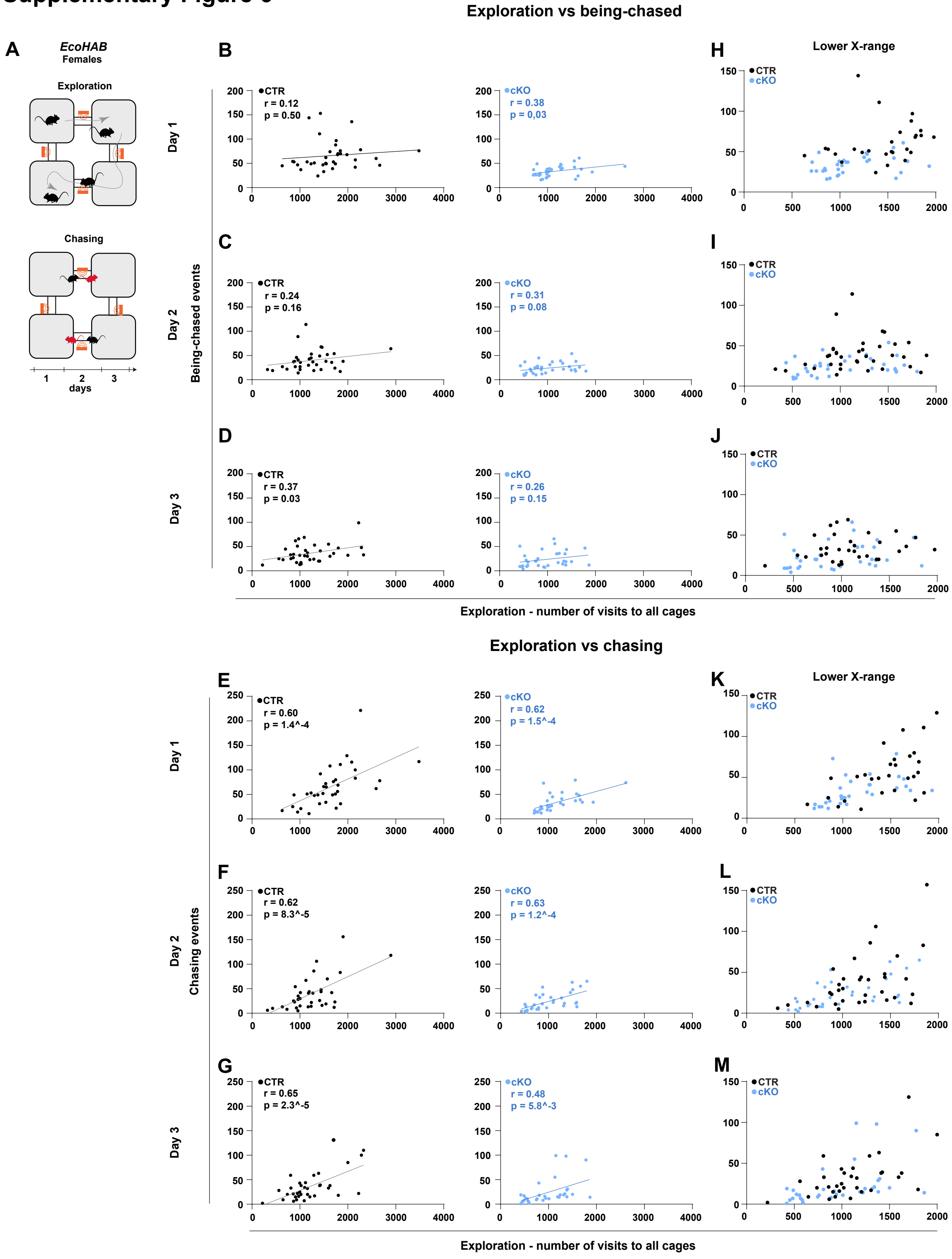

**SFig. 6. Dynamic social interactions in EcoHAB are not reflective of locomotor activity (mouse-by-mouse analysis)**

**(A)** Schematic of the EcoHAB setup, used to assess social behavior, illustrating two analyzed parameters: exploratory activity and chasing (black mice) or being chased (red mice). CTR n = 35 and cKO n = 32 in three independent cohorts for each genotype; only females were tested; results in B-G represent data from the dark phase. **(B–D)** Correlation analyses between exploratory activity and being chased in control (CTR) and *Cck*$^{Cre}$*:Tcf7l2*$^{fl/fl}$ (cKO) mice on days 1, 2 and 3. **(E-G)** Zoomed-in activity/being chased correlation plots restricted to the lower range of exploratory activity, showing CTR and cKO data plotted together for direct comparison. **(H-J)** Correlation analyses between exploratory activity and chasing on days 1, 2 and 3. **(K-M)** Zoomed-in activity/chasing correlation plots restricted to the lower range of exploratory activity, showing CTR and cKO data plotted together for direct comparison.

Dots represent individual mice. Data in B-D and E-J were analyzed using Pearson correlation test; r- and p-values are shown on the plots.

### Supplementary Figure 7

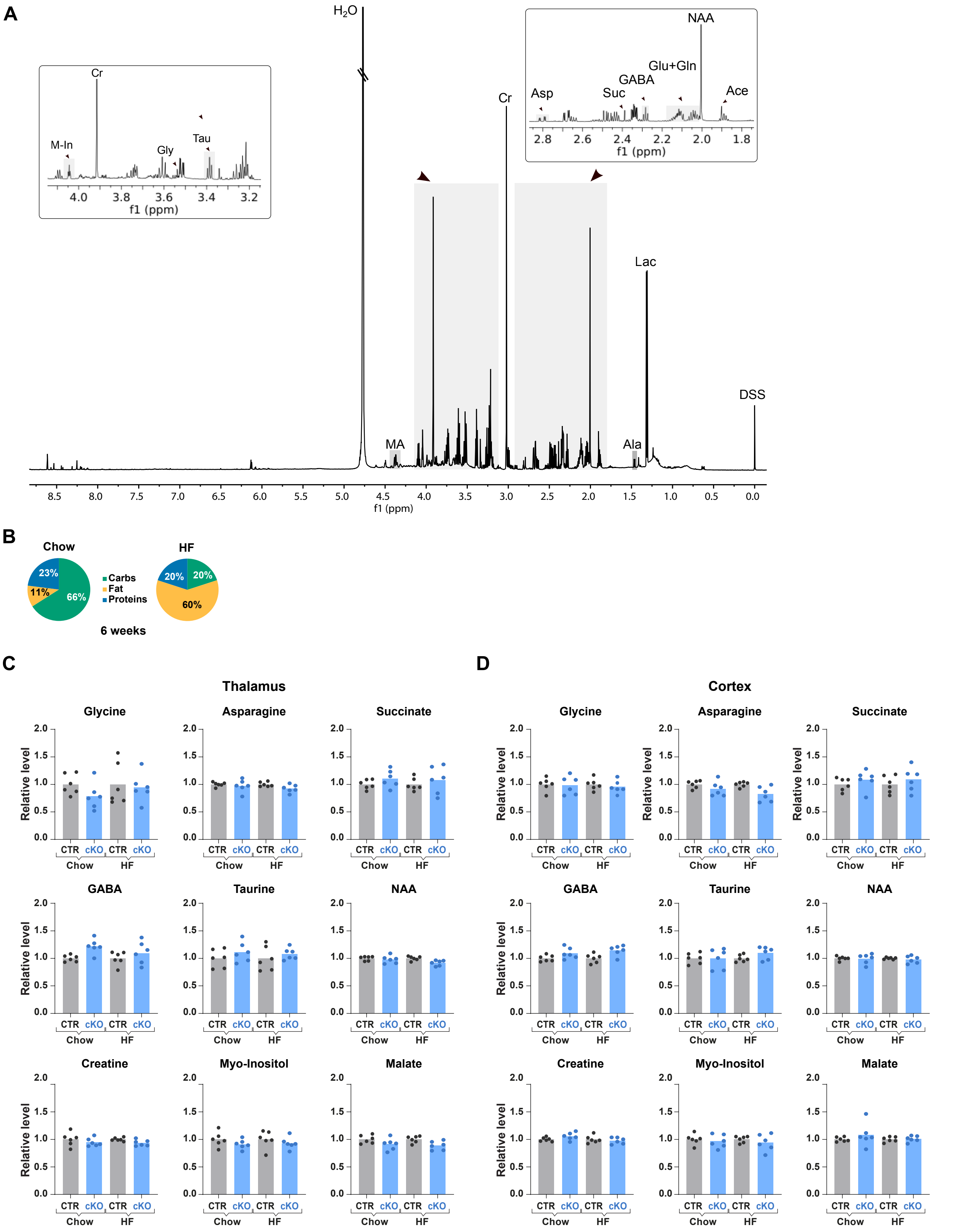

**SFig. 7. TCF7L2 depletion in the thalamus does not affect the levels of most of soluble metabolites in the brain**

**(A)** High-resolution 1H NMR spectrum (700MHz) of an aqueous extract of thalamic tissue. **(B)** Schematic of the dietary intervention showing the nutrient composition of the standard laboratory diet (Chow) and high-fat (HF) diet; only males were tested. **(C-D)** Levels of metabolites in the thalamus and cortex in control (CTR) and $Cck^{Cre}$:$Tcf7l2^{fl/fl}$ (cKO) male mice fed the chow or HF diet.
Bars in graphs represent mean values, and dots represent individual mice. Data in C-D were analyzed using one-way ANOVA followed by Tukey's multiple comparison test.

### Supplementary Figure 8

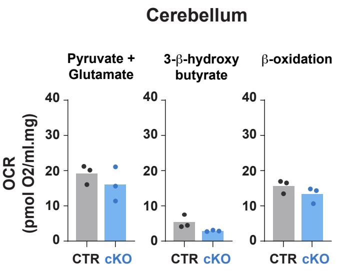


**SFig. 8. TCF7L2 depletion in the thalamus does not affect oxidation rates of metabolic substrates in the cerebellum**

Oxygen consumption rates (OCR) of permeabilized tissue membranes isolated from the cerebellum of control (CTR) and *Cck$^{Cre}$:Tcf7l2$^{fl/fl}$* (cKO) male mice. Data were obtained by sequentially adding fatty acids, 3-β-hydroxybutyrate, and pyruvate + glutamate to the Oxygraph-2k chamber.

Bars in graphs represent group means, and dots represent individual mice. Data were analyzed using an unpaired T-test.

#### Supplementary Figure 9

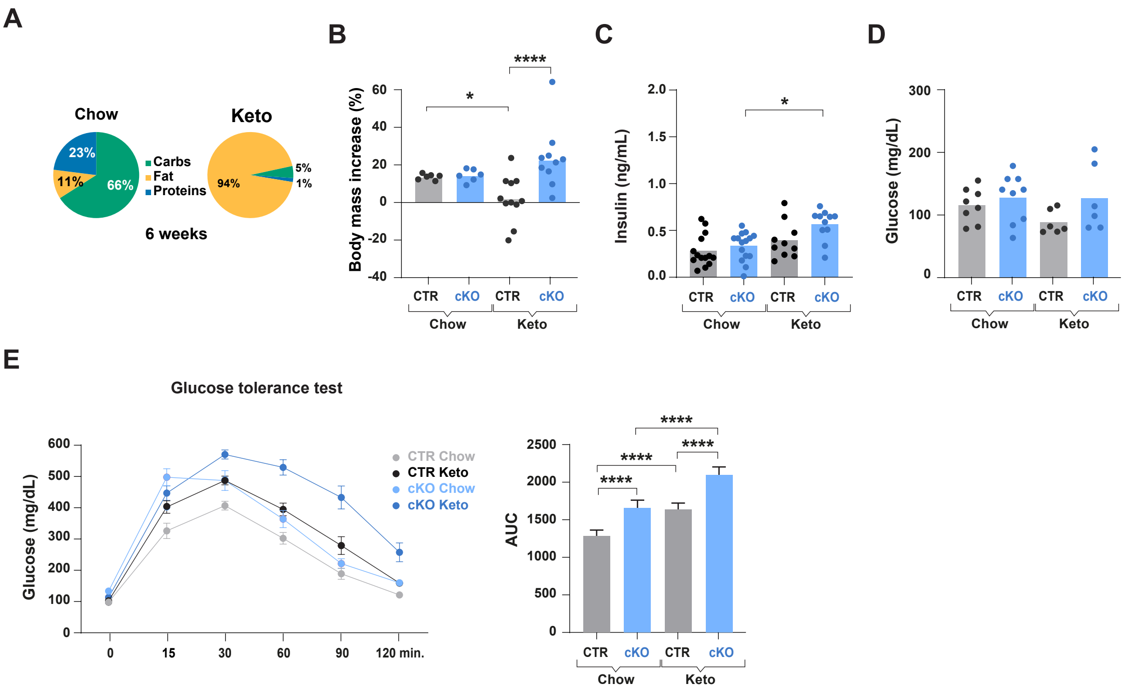


**SFig. 9. Ketogenic diet impairs systemic glucose tolerance in both control and *Tcf7l2-cKO* mice**

**(A)** Schematic of the dietary intervention showing the nutrient composition of the standard laboratory (Chow) and ketogenic (Keto) diets; only males were tested. For the Chow diet, the data from Fig. 1 were used. **(B)** Percentage increase in body mass for control (CTR) and *Cck*$^{Cre}$:*Tcf7l2*$^{fl/fl}$ (cKO) mice fed either the chow or ketogenic diet. **(C)** Fasting blood insulin levels. **(D)** Fasting blood glucose level. **(E)** Glucose tolerance test (GTT; left panel); and area under the curve (AUC) of GTT for the same groups (right panel); n = 6. Bars in graphs represent group means, dots in E left panel represent group means, and dots in B-D represent individual mice. Data in B-D, and E right panel were analysed using two-way ANOVA, followed by Tukey's multiple comparison test. *$p < 0.05$, ****$p < 0.0001$.

### Supplementary Figure 10

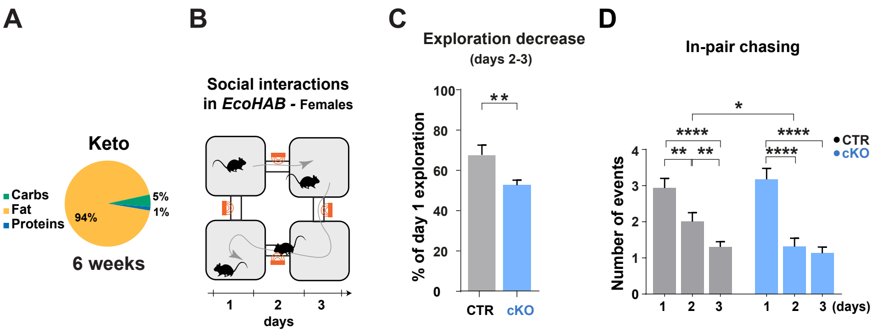


**SFig. 10. Ketogenic diet ameliorates social behaviour deficits induced by thalamic *Tcf7l2* knockout**

**(A)** Schematic of the dietary intervention showing the nutrient composition of the ketogenic (Keto) diet. **(B)** Schematic of the EcoHAB setup used to assess social behavior. CTR n = 21 and cKO n = 20 in two independent cohorts for each genotype; only females were tested; results in C-D represent data from the dark phase. **(C)** Percentage decrease in exploratory activity (visits to all cages) in days 2-3 relative to day 1 in control (CTR) and $Cck^{Cre}$:$Tcf7l2^{fl/fl}$ (cKO) mice fed the ketogenic diet. **(D)** In-pair chasing events, measured as one mouse trailing another through a tunnel for each mouse pair; the bar plot corresponds to the violin plot shown in Fig. 5I.

Bars in graphs represent mean values, with dots indicating individual mice. Data in C were analyzed using an unpaired T-test; data in D were analyzed using repeated measures two-way ANOVA, followed by Tukey's multiple comparison test. *$p < 0.05$, **$p < 0.01$, ***$p < 0.001$, ****$p < 0.0001$.
